# CD4^+^ T cells are homeostatic regulators during *Mtb* reinfection

**DOI:** 10.1101/2023.12.20.572669

**Authors:** Joshua D. Bromley, Sharie Keanne C. Ganchua, Sarah K. Nyquist, Pauline Maiello, Michael Chao, H. Jacob Borish, Mark Rodgers, Jaime Tomko, Kara Kracinovsky, Douaa Mugahid, Son Nguyen, Dennis Wang, Jacob M. Rosenberg, Edwin C. Klein, Hannah P. Gideon, Roisin Floyd-O’Sullivan, Bonnie Berger, Charles A Scanga, Philana Ling Lin, Sarah M. Fortune, Alex K. Shalek, JoAnne L. Flynn

## Abstract

Immunological priming – either in the context of prior infection or vaccination – elicits protective responses against subsequent *Mycobacterium tuberculosis* (*Mtb*) infection. However, the changes that occur in the lung cellular milieu post-primary *Mtb* infection and their contributions to protection upon reinfection remain poorly understood. Here, using clinical and microbiological endpoints in a non-human primate reinfection model, we demonstrate that prior *Mtb* infection elicits a long-lasting protective response against subsequent *Mtb* exposure and that the depletion of CD4^+^ T cells prior to *Mtb* rechallenge significantly abrogates this protection. Leveraging microbiologic, PET-CT, flow cytometric, and single-cell RNA-seq data from primary infection, reinfection, and reinfection-CD4^+^ T cell depleted granulomas, we identify differential cellular and microbial features of control. The data collectively demonstrate that the presence of CD4^+^ T cells in the setting of reinfection results in a reduced inflammatory lung milieu characterized by reprogrammed CD8^+^ T cell activity, reduced neutrophilia, and blunted type-1 immune signaling among myeloid cells, mitigating *Mtb* disease severity. These results open avenues for developing vaccines and therapeutics that not only target CD4^+^ and CD8^+^ T cells, but also modulate innate immune cells to limit *Mtb* disease.

## INTRODUCTION

Management of the tuberculosis (TB) pandemic is limited by the lack of a robust vaccine that protects against *Mycobacterium tuberculosis (Mtb*) infection and disease progression. Bacillus Calmette–Guérin (BCG) remains the only licensed TB vaccine. BCG offers protection against ∼70% of severe miliary and meningeal infections in pediatric TB; however, it fails to confer robust protection against infection or TB disease in adults (Lange et al., 2022). Regardless of BCG vaccination status, the majority (∼90%) of infected individuals can control *Mtb* bacilli naturally and experience asymptomatic infection (clinically classified as latent TB infection; LTBI). Only the minority (∼5-10%) experience overt clinical manifestations of disease (Lawn and Zumla, 2011). In TB endemic regions – where people are likely repeatedly exposed – the incidence rate of recurrent TB disease, either by relapse or reinfection, after successful treatment with antibiotics was 18 (China) and 14.6 (Spain) times higher than the incidence rate of initial TB disease in the general population (J-P Millet et al., 2009; Shen et al., 2017; Verver et al., 2005). However, the effects of prior *Mtb* infection appear to be contextual: a retrospective epidemiological meta-analyses of healthcare professionals reported a 79% lower risk of developing active TB in LTBI individuals after re-exposure to *Mtb* compared to uninfected individuals (Andrews et al., 2012). This observation is further bolstered by findings from non-human primate (NHP) and murine models, which demonstrate that concomitant immunity (immunological memory conferred by concurrent *Mtb* infection) provides robust protection against *Mtb* reinfection, and that this protection persists to some extent after drug treatment (Cadena et al., 2018; Ganchua et al., 2023; Nemeth et al., 2020).

Several factors could explain variation in a host’s ability to control TB following *Mtb* reinfection. Examples include intrinsic differences in host susceptibility, differences in the quality of the memory immune response, emergence of TB related structural lung disease, and pathogen characteristics (Abel et al., 2018; Cohen et al., 2022; Coscolla and Gagneux, 2014; Galagan, 2014). Since prior *Mtb* infection provides protection against reinfection in NHPs, we can use this model to dissect the roles that key immune cell subsets play in protection (Cadena et al., 2018). Here, we sought to define the roles and functions of CD4^+^ T cells in the setting of reinfection.

The importance of CD4^+^ T cells for protection from *Mtb* infection and TB disease has been established in humans by observing the devastating effects of HIV on TB disease burden worldwide. It is further supported by numerous studies in mice and NHPs, where loss of CD4^+^ T cells leads to increased pathology, bacterial burden, and reactivation of disease (Kwan and Ernst, 2011; Lin et al., 2012b; Scanga et al., 2000). However, in NHPs as in humans, the outcome of *Mtb* infection varies significantly across sites of infection, such that there can be simultaneous sterilization and progression of disease in different granulomas within the same host (Gideon et al., 2022; Lin et al., 2014). Likewise, while CD4^+^ T cell depletion in primary infection leads to worsened control on the whole, some lesions are fully sterilized and some animals do well – observations which the current paradigm of protective immunity to *Mtb* cannot explain (Diedrich et al., 2020; Foreman et al., 2022; Larson et al., 2023; Lin et al., 2012b).

In this study, we sought to evaluate systematically long-lived immunological reprogramming in pulmonary granulomas after primary and secondary infection and to elucidate the role of CD4^+^ T cells in protection against reinfection. Using *in vivo* perturbations (reinfection and CD4^+^ T cell depletion) and a combination of clinical, microbiologic, and high-dimensional single-cell transcriptomic analyses, we characterized intra- and inter-cellular changes associated with disease outcomes within pulmonary granuloma in cynomolgus macaques. Our analysis helps to unravel the intricacies of host-pathogen dynamics in TB, providing foundational insights for advancing vaccine research and therapeutic modalities.

## RESULTS

### Experimental design

This study aimed to determine the role of CD4^+^ T cells in establishing microbiologic, radiographic, and immunologic outcomes in the setting of *Mtb* reinfection in cynomolgus macaques. We used antibody-based depletion of CD4^+^ T cells (hereafter, αCD4) immediately prior to reinfection to assess CD4^+^ T cells effector functions rather than their roles in establishing adaptive responses to *Mtb* during primary infection. We compared the outcomes of *Mtb* reinfection in the setting of αCD4 (reinfection, CD4^+^ T cell depletion, n=7) with those in animals which received an isotype control (reinfection, IgG antibody infusion, n=6). We also examined primary infection in naïve animals (primary infection only, n=6). Overall, this enabled us to compare the outcome of reinfection to primary infection (IgG vs naïve), and then assess the impact of CD4^+^ T cell depletion (IgG vs αCD4) (**Figure 1A**).

**Figure 1.**
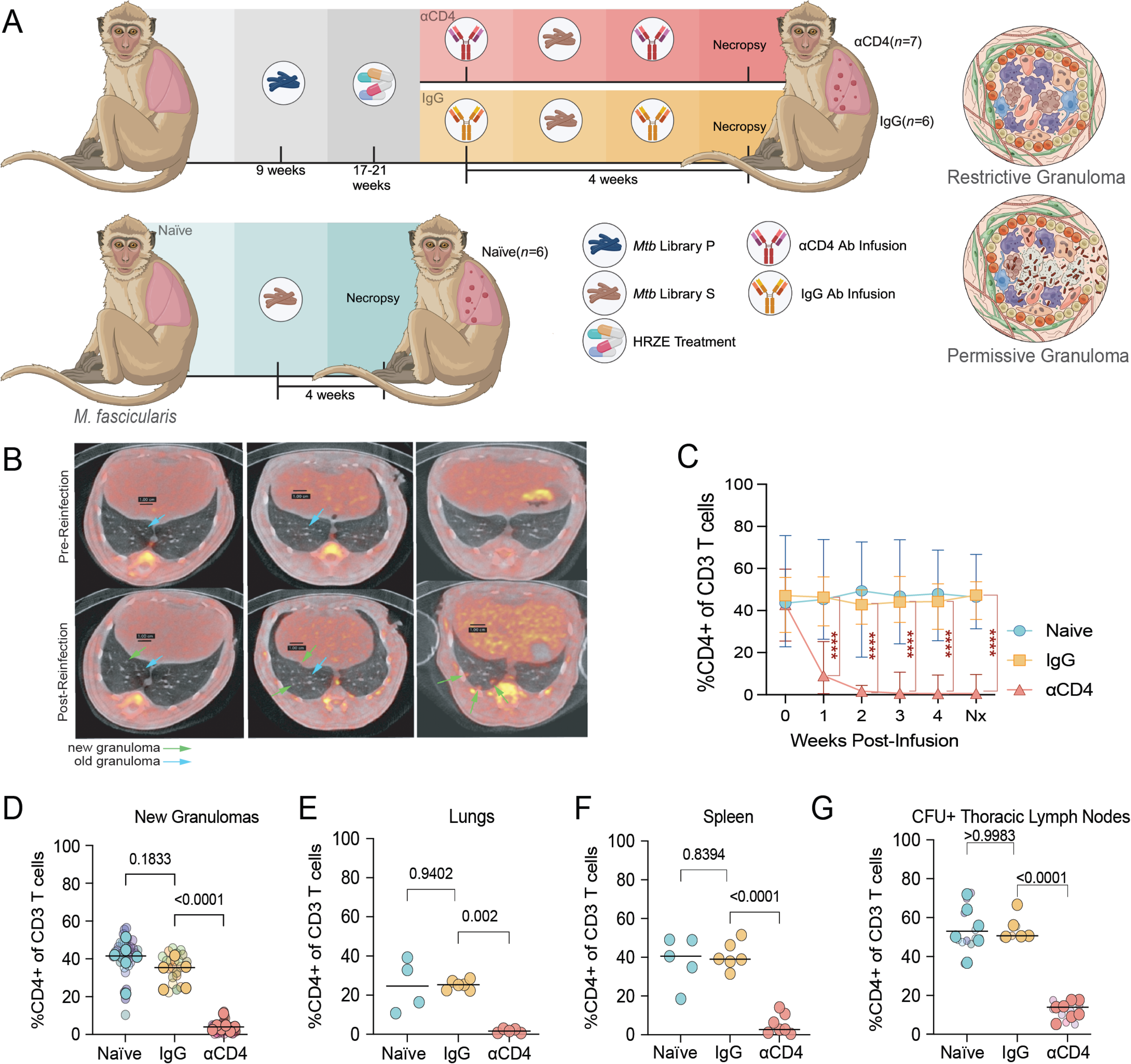
Experimental design. **(A)** Overview of cynomolgus macaque sample processing for clinical, microbiologic, and immunologic data (created with BioRender.com). **(B)** PET-CT scan of representative NHPs pre- and post-HRZE treatment. Old granulomas shown with blue arrows; new granulomas shown with green arrows. Left panel: IgG; middle panel: aCD4; right panel: naïve. **(C)** Fraction of CD3 expressing the cell surface marker CD4 post-antibody infusion in peripheral blood. Median and range shown (****, p<0.0001; mixed-effects model with Dunnett’s multiple comparisons test). **(D)** CD3^+^, CD4^+^ cells derived from TB granulomas. **(E)** CD3^+^, CD4^+^ cells from uninvolved lung tissue from *Mtb* infected macaques. **(F)** CD3^+^, CD4^+^ cells from the spleen of *Mtb* infected macaques **(G)** CD3^+^, CD4^+^ cells from CFU^+^ LNs of *Mtb* infected macaques. (D-G) Transparent smaller dots represent granulomas, colored by animal. Larger dots represent mean per animal and lines represent medians. One-way ANOVA with Dunnett’s multiple comparisons test.

We used *Mtb* Erdman for primary infection and reinfection. As previously described, these bacteria were differentially barcoded with a library of randomized tags, enabling us to track the infecting libraries (designated primary (P) and secondary (S)) and each unique founding bacterium within a library via deep sequencing (Martin et al., 2017). Specifically, the reinfection group was infected with ∼10 CFU of *Mtb* Library P, and infection was allowed to progress for nine weeks before the animals received the standard four-drug TB treatment regimen (isoniazid, rifampin, pyrazinamide, ethambutol; HRZE) for four to five months, with the duration of treatment guided by the resolution of disease as defined by PET-CT imaging. For this study, the animals were treated with drugs so that subsequent CD4 depletion would not result in overwhelming disease due to the primary infection. Our previous data support that drug treatment of primary infection has a modest effect on protection against reinfection (Ganchua et al., 2023). Subsequently, reinfection animals were randomized into the IgG or αCD4 cohorts. These groups began receiving weekly IgG or αCD4 antibody infusions starting one week before secondary infection with Library S and continuing until necropsy at four weeks post-infection. The naïve group was infected with ∼10 CFU of barcoded *Mtb*-Erdman (Library S) at the same time as the reinfection macaques, and infection progressed for four weeks before necropsy. Assessment of total lung FDG activity pre- and post-HRZE indicated that the responses to treatment were similar in each of the two antibody treatment groups (**Figure S1A**). PET-CT was continued over the course of the experiment, enabling the identification of newly formed granulomas following *Mtb* re-challenge and antibody infusions (**Figure 1B**). At necropsy, individual PET-CT scan-matched granulomas and lymph nodes, as well as all lung lobes, were resected and dissociated into single-cell suspensions for flow cytometric, scRNA-seq, and/or microbiologic analyses.

Over the course of *Mtb* reinfection and antibody infusion, peripheral blood was sampled weekly to assess CD4^+^ T cell depletion efficacy and to quantify cell type frequencies. CD4^+^ T cells were significantly depleted post-infusion (10- to 1,000-fold compared to pre-infusion levels) in the blood of αCD4 animals up until necropsy, while no changes in CD4^+^ T cell levels were observed in the naïve and IgG cohorts (**Figure 1C** and **S1B**). CD4^+^ T cell depletion in macaques also reduced the number of CD4^+^ T cells, but not CD8^+^ T cells or B cells in tissues, including granulomas and lymph nodes, as compared to macaques that received IgG Ab infusion (**Figure 1D-G** and **S1C-F**).

### Reinfection with *Mtb* reduces granuloma formation, as well as bacterial burden and dissemination in a CD4^+^ T cell dependent manner

Analysis of PET-CT scans after secondary infection showed that a similar number of granulomas formed in animals receiving IgG as compared to naïve and αCD4 treated animals, although there was a trend (p=0.0714) towards lower numbers of new granulomas in IgG treated compared to naïve animals (**Figure 2A**). This was in contrast to our previous study of reinfection in non-drug treated animals, where fewer granulomas were established in animals with a concurrent primary infection following reinfection (Cadena et al., 2018). Consistent with our prior data, granulomas in animals receiving IgG had significantly fewer viable bacteria than those in naïve animals, with CD4^+^ T cell depletion partially abrogating protection against reinfection (**Figure 2B, E-G** and **S2A)**. While there was a trend towards lower cumulative bacterial burdens (chromosomal equivalents (CEQ); an estimate of total live and dead bacilli) in granulomas from NHPs receiving IgG infusion, these differences did not reach statistical significance (**Figure 2C** and **S2B**). The same was true for the CFU:CEQ ratio – a proxy for bacterial killing within a granuloma (**Figure 2D** and **S2C,D;** Lin et al., 2014). These data are most consistent with a model in which previous infection leads to an immune environment that restricts bacterial growth in a CD4^+^ T cell dependent fashion but does not prevent establishment of infection or drive substantively increased bacterial killing.

**Figure 2.**
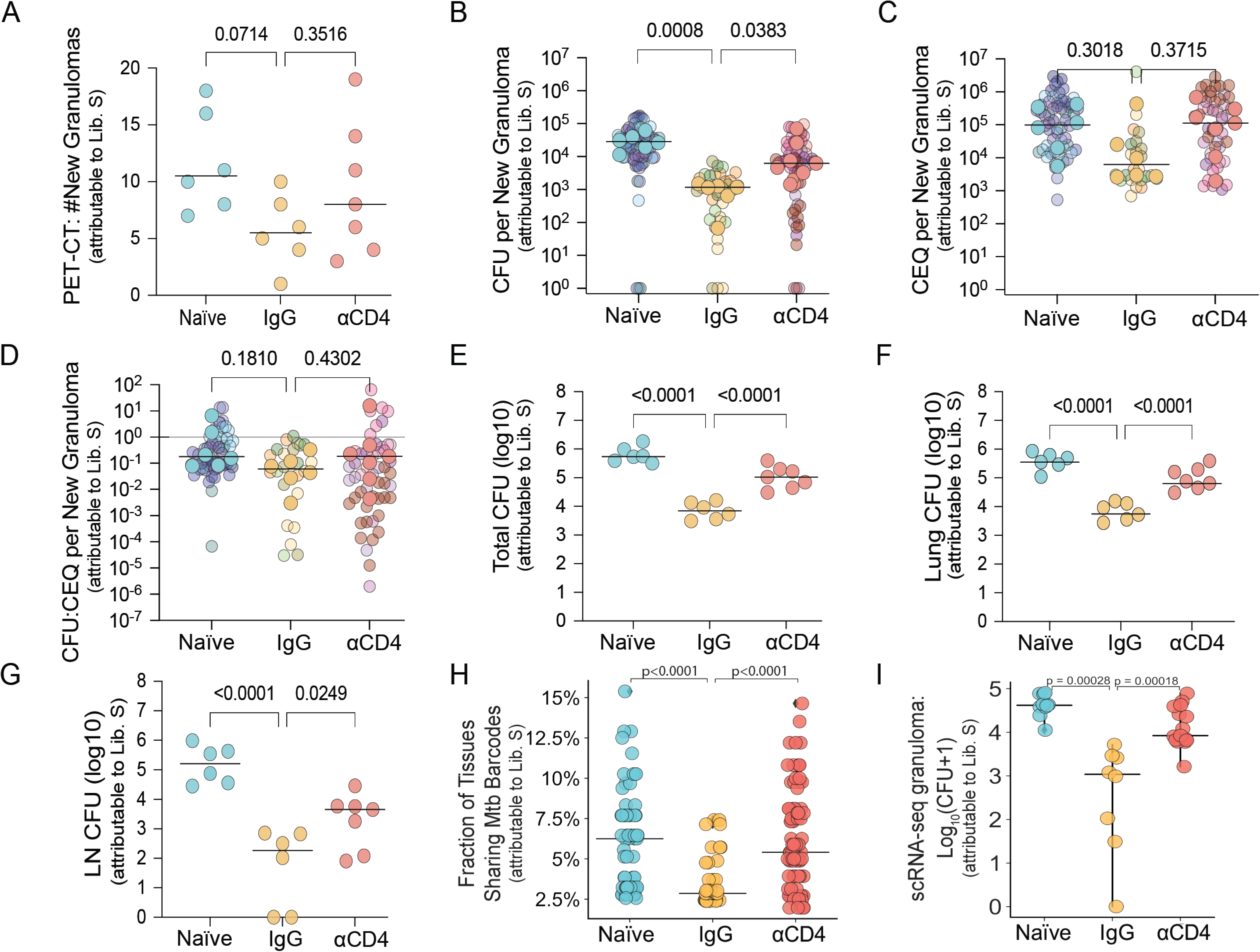
Reinfection with *Mtb* reduces granuloma formation, as well as bacterial burden and dissemination in a CD4^+^ T cell dependent manner. **(A)** Number of new granulomas identified using PET-CT following infection with *Mtb* library S. Lines represent medians. One-way ANOVA with Dunnett’s multiple comparison test, adjusted p-values reported. **(B)** Median number of viable *Mtb* colony forming units (CFU) per macaque (Kruskal-Wallis with Dunn’s multiple correction). Solid dots represent the median CFU per animal; lines represent medians. Transparent dots represent the median CFU of individual granulomas. **(C)** Median number of chromosomal equivalents (CEQ) per macaque (Kruskal-Wallis with Dunn’s multiple correction). Solid dots represent the median CFU per animal; lines represent medians. Transparent dots represent the median CFU of individual granulomas. **(D)** CFU:CEQ ratio, a proxy for bactericidal activity (Kruskal-Wallis with Dunn’s multiple correction). Solid dots represent the median CFU per animal; lines represent medians. Transparent dots represent the median CFU of individual granulomas. **(E)** Total CFU from granuloma, uninvolved lung tissue, and thoracic lymph nodes tissue. **(F)** Lung CFU from granuloma and uninvolved lung tissue. **(G)** Thoracic lymph node CFU. (E – G) Lines represent medians. One-way ANOVA with Dunnett’s multiple comparisons test. **(H)** Individual granuloma *Mtb* CFU. Individual dots represent single granuloma subject to Seq-Well S^3^ scRNA-seq (Mann-Whitney U Test). **(I)** Fraction of tissues (lymph node, spleen, lung) sharing library S barcodes (Kruskal Wallis with Dunn’s multiple comparison test).

*Mtb* barcode analysis of samples retrieved at necropsy revealed that previous infection provided enhanced protection against *Mtb* dissemination to lymph nodes and this protection was partially dependent on CD4^+^ T cells (**Figure S2E-G**). We enumerated barcode dissemination and found a lower percentage of shared bacterial barcodes between tissues in the IgG animals as compared to both naïve and αCD4-treated animals, further suggesting that reinfection reduces bacterial dissemination in a CD4^+^ T cell dependent manner (IgG vs naïve, p<0.0001; IgG vs αCD4, p<0.0001, Kruskal Wallis with Dunn’s multiple comparison test; **Figure 2I**).

### Cellular remodeling of the TB granuloma microenvironment following *Mtb* reinfection

To define the cellular features associated with protection in the setting of reinfection and the effects of CD4^+^ T cell depletion, we performed Seq-Well S^3^-based massively-parallel scRNA-seq on granulomas isolated from the three experimental groups (**Figures 2H, 3A;** Hughes et al., 2020). We analyzed 33 granulomas that were confirmed to arise from the second infection (Library S) (naïve=10, IgG=8, αCD4=15) from 7 cynomolgus macaques (naïve=2, IgG=3, αCD4=3), yielding a total of 88,360 high-quality transcriptomes. We annotated 16 clusters corresponding to distinct immune and non-immune cell types based on known marker genes and reference signatures (**Figure 3A** and **S3A;** Sikkema et al., 2023). While cellular frequencies varied among individual granulomas and experimental groups, each cluster was represented by multiple samples (**Figure S3B**).

**Figure 3.**
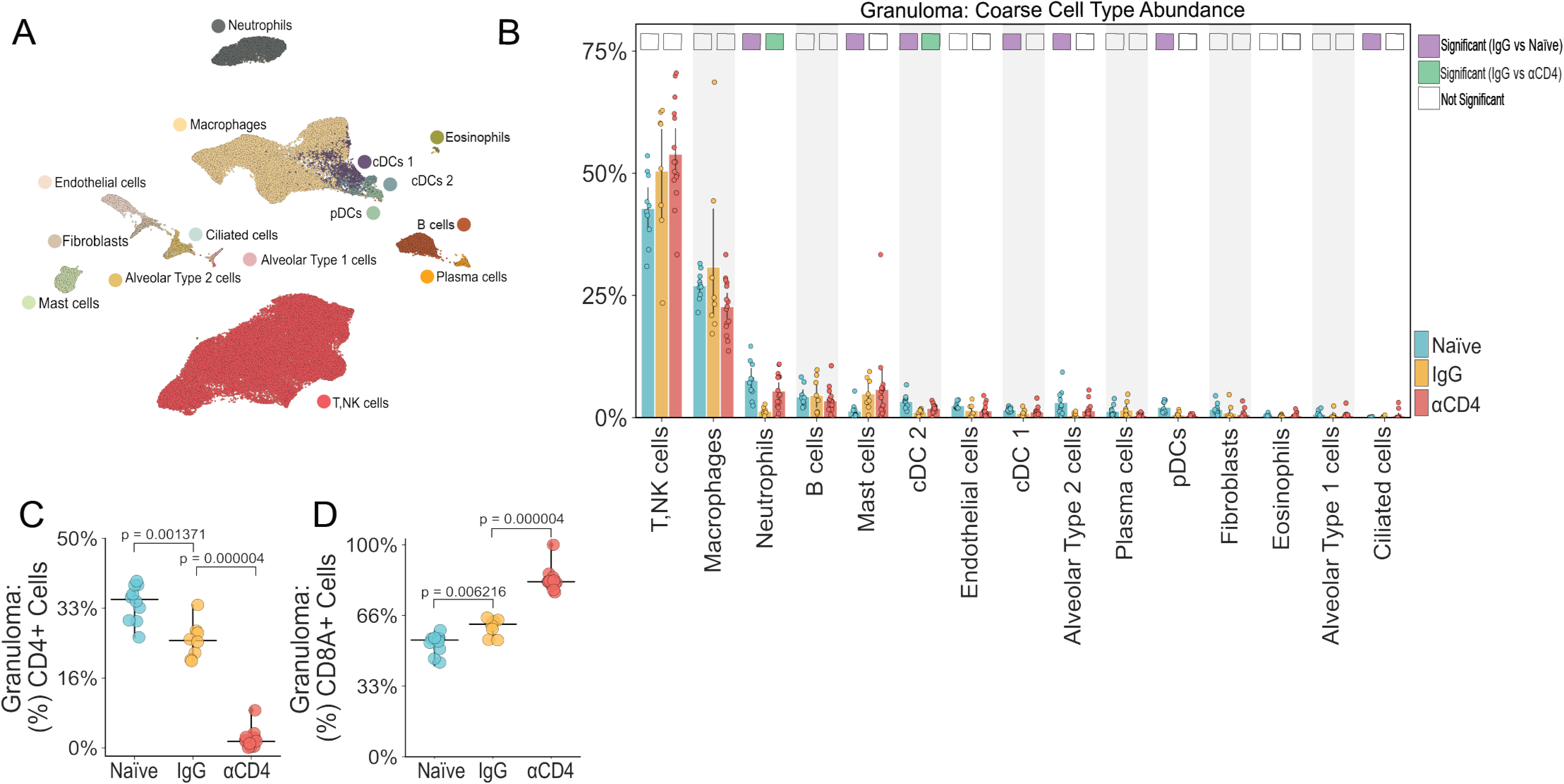
Cellular remodeling of the TB granuloma microenvironment following *Mtb* reinfection. **(A)** UMAP embedding of Seq-Well S^3^ derived granuloma transcriptomes colored by coarse cell type. **(B)** Coarse cell type frequencies colored by experimental group. Individual dots represent single granuloma. Differentially abundant IgG vs naïve (purple) and IgG vs αCD4 (green) marked with colored square. Cell types are differentially abundant if significant using two of three methods: Mann-Whitney U test; scCODA Bayesian model, and Fishers exact test. **(C)** Fraction of granuloma T, NK cells expressing *CD4* from Seq-Well S^3^ derived transcriptomes (Mann-Whitney U Test). **(D)** Fraction of granuloma T, NK cells expressing *CD8A* from Seq-Well S^3^ derived transcriptomes (Mann-Whitney U Test).

We next sought to identify whether there were significant changes in cell type frequencies across granulomas. We implemented multivariate (scCODA), univariate (Mann-Whitney U test), and nonparametric (Fisher’s exact test) tests to account for perturbation, cell type codependences, and low sample size, and considered a cell type differentially abundant if significant by at least two tests (Büttner et al., 2021; Smillie et al., 2019). We first assessed the global T cell composition of lesions from the three groups and observed a trend toward higher overall T, NK cell frequencies among IgG granulomas relative to naïve (**Figure 3B**). Given the trend toward increased T, NK cell frequencies among lesions formed in the setting of reinfection relative to naïve granulomas even in the setting of CD4^+^ T cell depletion, we sought to identify whether prior infection promotes CD4^+^ or CD8^+^ T cell recruitment to the granuloma. Surprisingly, the total fraction of *CD4*+ and *CD8+* T cells among *CD3D* expressors was significantly lower and higher in IgG granulomas relative to naïve granulomas, respectively (p=0.0014 and p=0.0062, respectively, Mann-Whitney U test, **Figure 3C,D**). The former is in line with flow cytometric data demonstrating a lower frequency of CD4+ T cells in IgG granulomas, relative to naïve granulomas (**Figure 1D**). There was no difference in the frequency of total T, NK cells between IgG and αCD4 granulomas, but the latter had significantly fewer *CD4*^+^ T cells and significantly more *CD8^+^*T cells (**Figure 3C,D**).

IgG granulomas had lower frequencies of neutrophils relative to naïve granulomas, and depletion of CD4^+^ T cells led to increased frequencies of neutrophils. IgG granulomas also had lower frequencies of cDC2s relative to both naïve and αCD4 granulomas. Finally, there were several differences between naïve and IgG granulomas that were not affected by CD4^+^ T cell depletion. Mast cells were more frequent in IgG lesions compared to naïve granuloma, but this was not altered by CD4^+^ T cell depletion. IgG granulomas also had lower frequencies of cDC1s, alveolar type 2 cells, ciliated cells, and pDCs as compared to naïve granulomas.

### CD4^+^ T cells regulate immune tone in granulomas formed after reinfection with *Mtb*

To better understand how reinfection and CD4^+^ T cell depletion influence the structure of the overall T, NK cell cluster, we next classified all *CD3D,E, CD4, CD8A,B* T, NK cells into 11 major subpopulations based on gene signatures from external single-cell datasets (**Figure 4A** and **S3C** and **Table 1**; Almanzar et al., 2020; Zheng et al., 2021). The proportions of several of these T, NK cell subsets significantly differed among reinfection (IgG) granulomas, particularly the *CD8* enriched (GZMK^hi^ T_EM/PEX-like_) and *CD8, CD4* co-expressing (T_EMRA-like_) subsets. We found significant enrichment of immuno-modulatory and -regulatory *CD8* GZMK^hi^ T_EM/PEX-like_ (*GZMK, EOMES, TOX, TIGIT, IL10*) (Mogilenko et al., 2021) cells in IgG granulomas relative to naïve ones; CD4-depletion significantly impaired either the localization or retention of these cells, suggesting CD4-dependence even after immune priming (**Figure 4B**).

**Figure 4.**
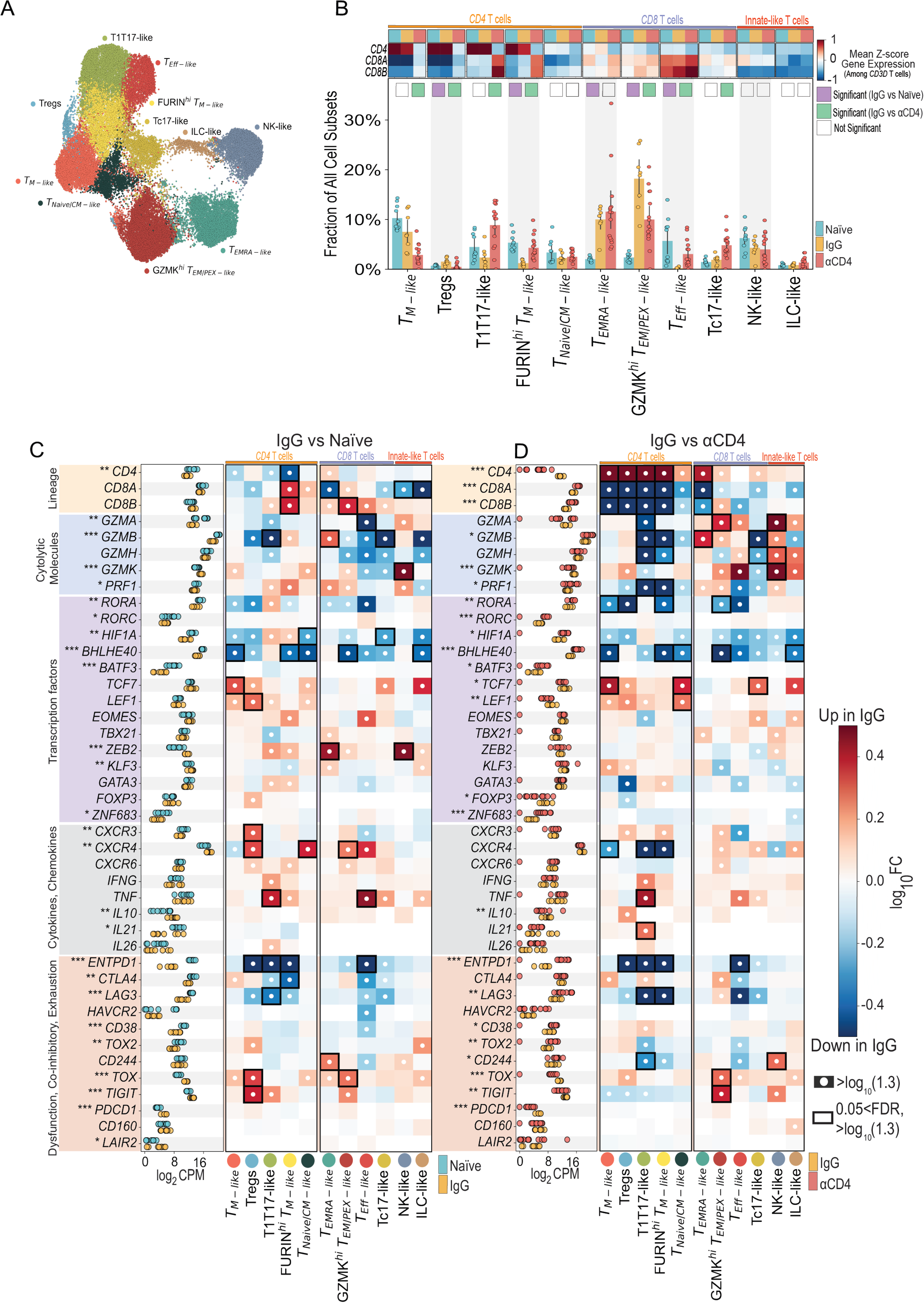
CD4^+^ T cells regulate T cell cellularity, cytokine flux, and immune tone in in the TB granulomas following *Mtb* reinfection. **(A)** UMAP embedding depicting T, NK cell subpopulations identified by sub-clustering. **(B)** Heatmap depicts gene expression levels (mean z-score) of T cell lineage markers *CD4, CD8A,* and *CD8B*. Columns represent gene expression in individual NHP groups – Naïve (light blue), IgG (yellow), αCD4 (red). Bar plot of T, NK subpopulation frequencies among all granuloma cellular subpopulations colored by experimental group. Differentially abundant IgG vs naïve (purple) and IgG vs αCD4 (green) marked with colored square. Cell types are differentially abundant if significant using two of three methods: Mann-Whitney U test; scCODA Bayesian model, and Fishers exact test. **(C-D)** T, NK cell pseudobulk Log_2_CPM for naïve (light blue), IgG (yellow), and αCD4 (red) NHP granulomas (*** p<0.001,** p<0.01, *p<0.05; Wilcoxon rank-sum test). Heatmap depicting log_10_FC of lineage markers, cytolytic molecules, select transcription factors, immunoregulatory molecules, and chemokines, and cytokines (rows) for each cell type (columns) in NHP granulomas IgG vs naïve (C) or IgG vs αCD4 lesions (D). White circles indicate >log_10_|1.3| fold change, relative to naïve or αCD4 granulomas. Black rectangles indicate 0.05 FDR and >log_10_|1.3| fold change, relative to naïve or αCD4 granulomas.

**TABLE 1.**
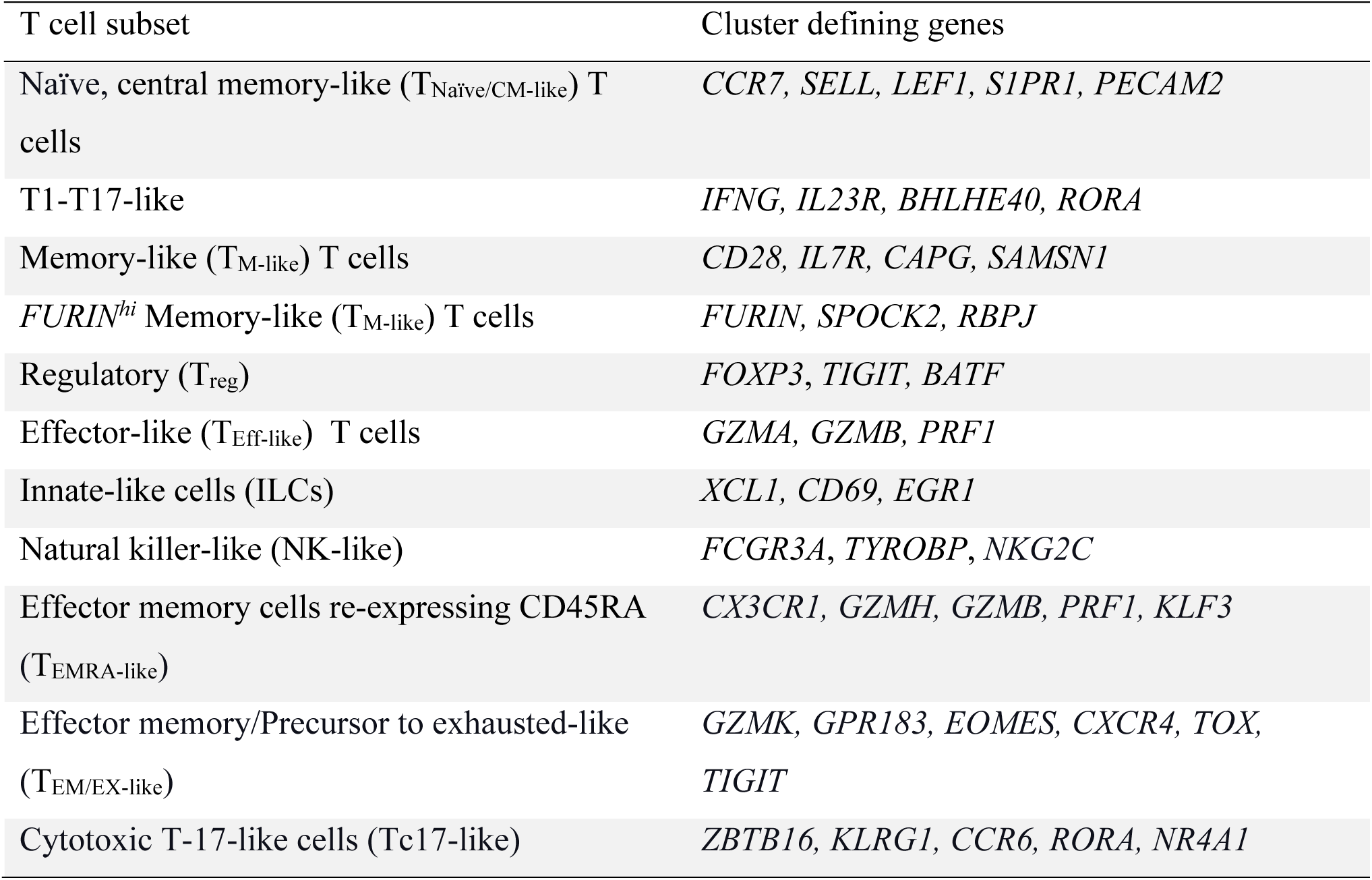

We next investigated how infection status and immune perturbation altered global T, NK cell responses, as well as those of each cell subpopulation, by performing pairwise (IgG vs naïve; IgG vs αCD4) T, NK pseudobulk (i.e., all T, NK subsets aggregated) differential gene expression (DGE), and pairwise DGE analyses within T cell subpopulations. Pairwise analyses among the 11 identified T, NK cell subsets revealed a total of 1,542 DE genes (773 upregulated, 759 downregulated) in IgG vs naïve lesions and 1,263 DE genes (783 upregulated, 480 downregulated) in IgG vs αCD4 granuloma, demonstrating significant shifts in T cell circuitry in immunologically primed animals relative to naïve (IgG vs naïve) or CD4-depletion (IgG vs αCD4) animals (**Figure 4C,D** and **S3F**).

To examine the potential functional significance of these DE genes, we systematically queried cytokines canonically associated with protective anti-*Mtb* responses, including *TNF*, *IFNG*, and the pleiotropic cytokine *IL10*, while excluding *IL17* due to its low expression (**Figure 4C**). There were no significant differences in global (pseudobulk) *TNF* or *IFNG* expression among IgG vs naïve granulomas. We similarly found no significant differences in *IFNG* expression among T, NK subsets. We did observe significant induction of *IL10* expression in IgG compared to naïve granulomas, globally (**Figure 4C**). However, there was no one T, NK subset that was significantly enriched for *IL10* expression among IgG lesions; rather, *IL10* was expressed across several subsets. In addition to higher expression of *IL10*, IgG granulomas were characterized by greater global expression of immunoregulatory (*TIGIT*, *TOX*), costimulatory (*CD2*, *CD28*, *ICOS*), and negative regulators (*CD5*, *CD6*) of T cell activation (**Figure S3D**). Notably, the elevated expression of *TIGIT* and *TOX* was primarily associated with Treg and GZMK^hi^ T_EM/PEX-like_ subsets. Relative to naïve granulomas, IgG granulomas had lower expression of most cytotoxic effector (*GZMA, GZMB, GZMH, PRF1*), hypoxia-induced factors (*HIF1A*, *BHLHE40*, *ENTPD1*), T1T17 transcription factors (TFs) (*RORA*, *RORC*) and interferon-stimulated genes (*ISG15, ISG20, IFI27*), across several T, NK subsets.

In the setting of CD4^+^ T cell depletion, there was a global reduction in the abundance of *IL10*, *PDCD1*, *TIGIT,* and *TOX* gene expression (**Figure 4D**). In αCD4 vs IgG granulomas, there was higher expression levels of cytolytic effector molecules (*GZMA, GZMB, GZMH, PRF1*) and T1T17-associated TFs. There was also increased expression of the PD-1-repressor *SATB1* and *BHLHE40*, a putative negative regulator of *IL10* expression and hypoxia-induced factor (**Figure S3E**; Huynh et al., 2018; Stephen et al., 2017). Collectively, these changes suggest CD8^+^ T cell reprogramming following *Mtb* reinfection and that acquisition of aspects of these terminally differentiated and immunoregulatory *CD8^+^* T, NK cell gene programs are CD4-dependent.

### Attenuated type-1 immunity among monocyte-derived transcriptomes in *Mtb* reinfection granulomas

Considering the established paradigm where CD4^+^ T cells orchestrate pro-inflammatory myeloid cell responses primarily through IFN-γ and TNF-mediated pathways, we explored whether the observed increase in immunoregulatory T cell phenotypes among IgG granulomas was associated with altered myeloid cellularity and transcriptional programming in reinfection. Monocyte-derived cells partitioned into 6 subpopulations and exhibited varying degrees of expansion or contraction in naïve, IgG, and αCD4 granulomas (**Figure 5A-B, S4A**).

**Figure 5.**
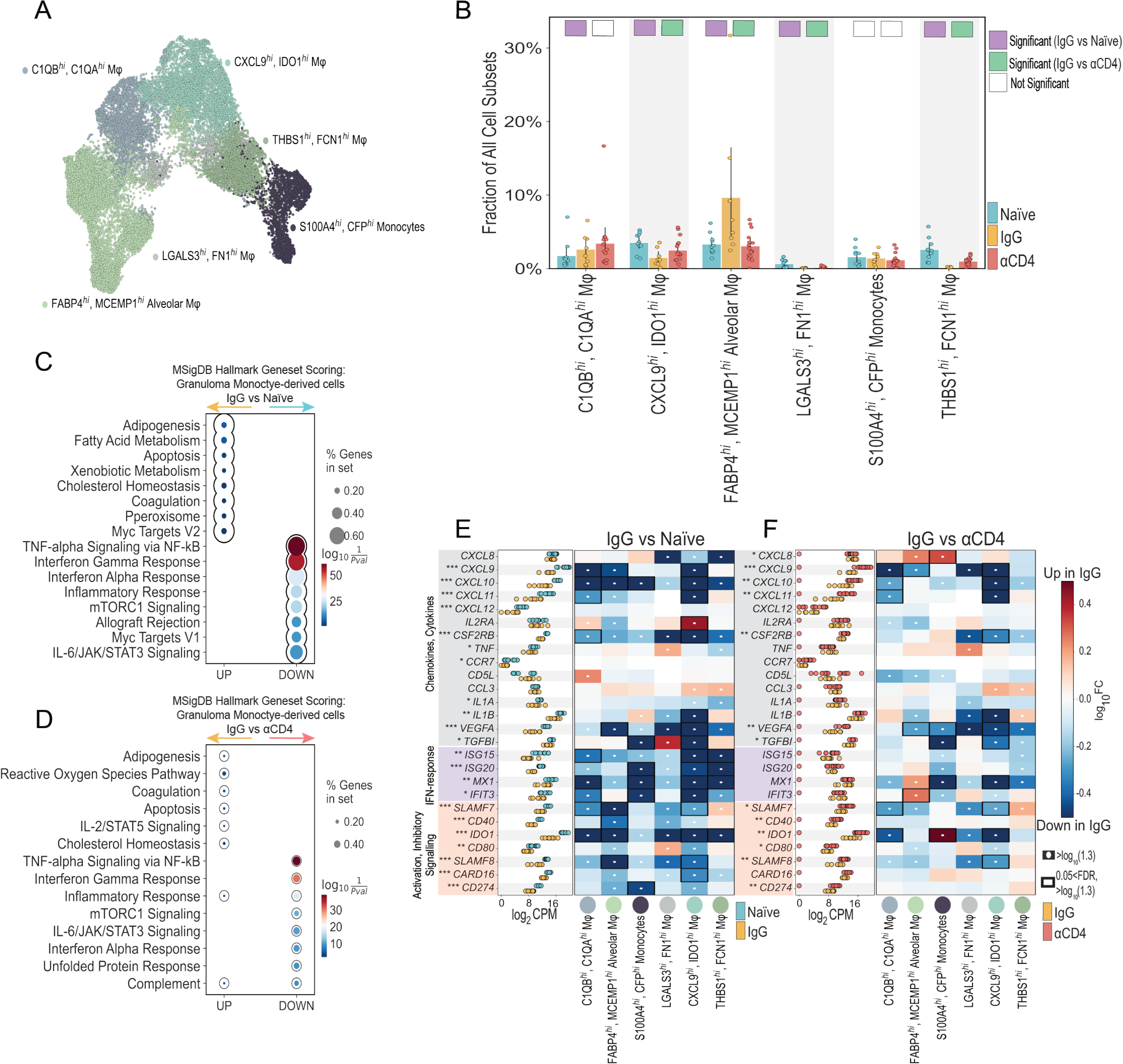
Attenuated type-1 immunity among monocyte-derived transcriptomes in *Mtb* reinfection granulomas. **(A)** UMAP embedding depicting Monocyte-derived cell identified by sub-clustering. UMAP embeddings depicting Monocyte-derived cell subpopulation densities, split by NHP cohort **(B)** Bar plot of Monocyte-derived progenitor frequencies, among all granuloma cell subpopulations, colored by experimental group. Individual dots represent single granuloma. Differentially abundant IgG vs naïve (purple) and IgG vs αCD4 (green) marked with colored square. Cell types are differentially abundant if significant using two of three methods: Mann-Whitney U test; scCODA Bayesian model, and Fishers exact test. **(C, D)** Enriched pathways from identified using differentially expressed genes (Mann-Whitney U test (Wilcoxon rank-sum) (p value<0.05)) from naïve, IgG, and αCD4 Sizes. Circles size represent to the number of genes in Hallmark Geneset, and color (Red-Blue) represents the geneset enrichment score. Genesets that are “up” (x-axis) are enriched among IgG granulomas, whereas “down” genesets are enriched among naïve **(C)** and αCD4 **(D)** granulomas, respectively. **(E-F)** Monocyte-derived pseudobulk Log_2_CPM for naïve (light blue), IgG (yellow), and αCD4 (red) NHP granulomas (*** p<0.001, ** p<0.01, *p<0.05; Wilcoxn rank-sum test). Heatmap depicting log_10_FC of select transcription factors, immunoregulatory molecules, and chemokines, and cytokines (rows) for each cell type (columns) in NHP granulomas IgG vs naïve (E) or IgG vs αCD4 lesions (F). White circles indicate >log_10_|1.3| fold change, relative to naïve or αCD4 granulomas. Black rectangles indicate 0.05 FDR and >log_10_|1.3| fold change, relative to naïve or αCD4 granulomas.

To identify global changes in myeloid gene programming following *Mtb* reinfection, we scored all monocyte-derived cells against Hallmark gene sets (**Figure 5C**). Specifically, all monocyte and macrophage subsets were grouped as “macrophages” and subject to gene set enrichment scoring. Our analyses demonstrated a global reduction in inflammatory responses, specifically IFNα-, IL6/JAK/STAT3, and IFNγ-responses, as well as significant enrichment of adipogenesis, fatty acid metabolism, Myc targets, and DNA repair signatures among monocyte-derived subsets in IgG relative to naïve granulomas. In contrast, in CD4-depletion granulomas the macrophages had increased IFNα-, IL6/JAK/STAT3, and IFNγ-inflammatory responses relative to IgG granulomas (**Figure 5D**).

Pairwise DGE analyses among monocyte-derived subpopulations revealed a total of 2,210 DE genes (1,236 upregulated, 974 downregulated) in IgG vs naïve lesions and 1,234 DE genes (777 upregulated, 457 downregulated) in IgG vs αCD4 granuloma (**Figure S4B**). Myeloid cells from naïve granulomas featured both significant global and subpopulation-specific increases in the expression of interferon-response genes (*ISG15*, *IRF7*), pro-inflammatory mediators (*IL1A*, *ILB*), chemokines and cytokines including the CXCR3 ligands (*CXCL9, CXLC10, CXCL11*), fibrosis-related genes (*VEGFA*, *TGFB1*), and immunoregulatory molecules (*IDO1, CD274* (PD-L1), *IL10*) relative to macrophages in IgG granulomas (**Figure 5E-F** and **S4C)**. In the absence of CD4^+^ T cells, a pseudobulk analysis indicated increased type-1 (e.g., *CXCL9-11*) immune signaling. A subpopulation-specific DE analysis of IgG vs αCD4 revealed various monocyte subsets significantly upregulated expression of CXCR3 ligands (*CXCL9, CXLC10, CXCL11*), and to a lesser featured upregulation of *IDO1* and *CD274* but not *IL10* in the αCD4 granulomas (**Figure 5F** and **S4D**).

Overall, naïve lesions showed enhanced type-1 immune signaling, while IgG granulomas displayed a significant reduction in these responses. The reversion of lesions toward a naïve-like state with CD4^+^ T cell depletion indicates a regulatory role for CD4^+^ T cells over the myeloid-driven inflammatory response during *Mtb* reinfection.

### Neutrophil heterogeneity in the TB granuloma

Neutrophils play a crucial role as frontline defenders against microbial infections and are quickly recruited to sites of inflammation upon *Mtb* infection. However, their role in TB disease remains enigmatic, as they promote both *Mtb* sterilization and pathology (Ravesloot-Chávez et al., 2021). To evaluate how prior infection modulates neutrophil recruitment and phenotype upon reinfection and the role of CD4^+^ T cells in that modulation, we quantified differences in cellular frequencies and gene expression following sub-clustering. Our analysis identified two neutrophil subpopulations: *ICAM1*^hi^, *NBN*^hi^ neutrophils (*ICAM1*, *CD274*, *GADD45B, CCL3L1*) and *SORL1*^hi^, *CFD^hi^* neutrophils (*SORL1*, *CFD*, *CORO1A*, *PLBD1*) (**Figure 6A** and **S4E**; Montaldo et al., 2022). Both neutrophil subpopulations were significantly underrepresented among IgG lesions compared to naïve and αCD4 granulomas, suggesting that either bacterial burden and/or CD4^+^ T cells regulate neutrophilic response and infiltration (**Figure 6B**).

**Figure 6.**
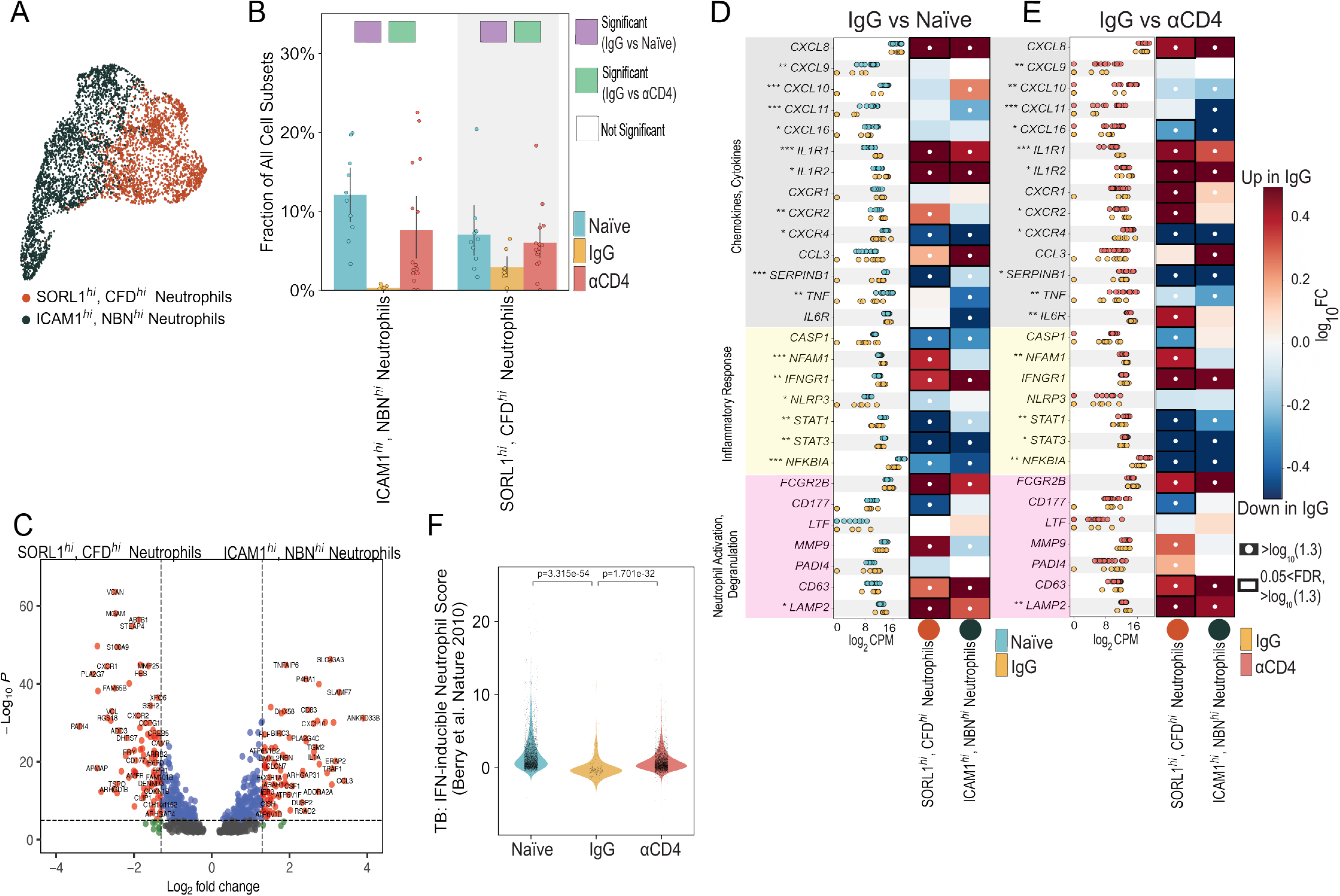
Neutrophil heterogeneity in the TB granuloma. **(A)** UMAP embedding depicting neutrophil cell subpopulations identified by sub-clustering **(B)** Bar plot of neutrophil subset frequencies, among all granuloma cell subpopulations, colored by experimental group. Individual dots represent single granuloma. Differentially abundant IgG vs naïve (purple) and IgG vs αCD4 (green) marked with colored square. Cell types are differentially abundant if significant using two of three methods: Mann-Whitney U test; scCODA Bayesian model, and Fishers exact test. **(C)** Volcano plot depicting pseudobulk differential gene expression (DESeq2) *ICAM1*^hi^, *NBN*^hi^ vs , *SORL1*^hi^, *CFD^hi^* neutrophils (for all NHP experimental groups). Volcano plot x-axis indicates the log_2_FC and y-axis indicates the -log10(pvalue). Vertical dashed lines represent log_2_FC threshold >|1.3|. Horizontal line indicates -log10(pvalue)>|0.05| threshold. **(D-E)** Neutrophil pseudobulk Log_2_CPM for naïve (light blue), IgG (yellow), and αCD4 (red) NHP granulomas (*** p<0.001,** p<0.01, *p<0.05; Wilcoxon rank-sum test). Heatmap depicting log_10_FC of select transcription factors, immunoregulatory molecules, and chemokines, and cytokines (rows) for each cell type (columns) in NHP granulomas IgG vs naïve (D) or IgG vs αCD4 lesions (E). White circles indicate >log_10_|1.3| fold change, relative to naïve or αCD4 granuloma. Black rectangles indicate 0.05 FDR and >log_10_|1.3| fold change, relative to naïve or αCD4 granulomas. **(F)** Violin plots of IFN-inducible neutrophil module scores, split by NHP group. Significance by Mann-Whitney U test.

To uncover potential differences in neutrophil transcriptional programming, we performed pairwise pseudobulk DGE analysis between these neutrophil subsets (*ICAM1*^hi^, *NBN*^hi^ vs *SORL1*^hi^, *CFD^hi^* neutrophils) and pairwise DGE analyses across conditions (naïve vs IgG, and αCD4 vs IgG) between these neutrophil subsets. Comparisons of *ICAM1*^hi^, *NBN*^hi^ to *SORL1*^hi^, *CFD^hi^* transcriptomes revealed *ICAM1*^hi^, *NBN*^hi^ neutrophils upregulate type-1 immune chemokines (*CXCL10*, *CXCL11*) and cytokines (*CCL3*, *IL1A*), whereas *SORL1*^hi^, *CFD^hi^* neutrophils upregulated molecules implicated in neutrophil trafficking (*CXCR1*, *CXCR2*) and netosis (*MGAM*, *MMP25*) (**Figure 6C**; Carmona-Rivera et al., 2015; Gasperini et al., 1999; Xie et al., 2020). Pairwise DE analyses (naïve vs IgG, and aCD4 vs IgG) revealed few significant differences in gene programming among *ICAM1*^hi^, *NBN*^hi^ transcriptomes; however, *SORL1*^hi^, *CFD^hi^* neutrophils had significantly altered transcriptomes among IgG lesions relative to naïve, with naïve granulomas expressing significantly higher levels of inflammatory response genes (*STAT1, STAT3, CASP1)*, type-1 immune chemokines (*CXCL10*, *CXCL11*) and cytokines (*CCL3L1*, *TNF*, *IL1A*) (Ichikawa et al., 2013). CD4 depleted lesions, meanwhile, exhibited similar ‘naïve-like’ neutrophil gene programming compared to IgG lesions (**Figure 6D-E** and **S4F**). We further scored neutrophils against an IFN-inducible neutrophil gene signature previously shown to be upregulated in humans with active TB (**Figure 6F**; Berry et al., 2010). In line with our DE analyses, IgG neutrophils featured significant blunting of IFN-inducible genes.

Collectively, our data delineate the diversity among granuloma-localized neutrophils and demonstrate a significant reduction in neutrophilic responses among IgG lesions compared to naïve or αCD4, implying a potential regulatory role for CD4^+^ T cells on neutrophil-driven immunity, pathophysiology, and TB disease progression. Furthermore, these data support the model that increased neutrophilic infiltration may contribute to the formation of *Mtb-*permissive niches, thus contributing to elevated bacillary loads among naïve and αCD4 lesions (Lovewell et al., 2021).

### Differential cell-cell interactions in immunologically primed granulomas

To assess how the different aforementioned factors act together to modulate host immunity, we investigated differential cell-cell interaction networks among naïve, IgG, and αCD4 granulomas (i.e., IgG vs naïve, and IgG vs αCD4; Browaeys et al., 2023). First, we identified differences among coarse-level cell-cell interactions occurring in primary infection granulomas (naïve) vs those formed in a primed immune environment (IgG). Relative to IgG lesions, naïve cell-cell interaction networks were dominated by signaling from neutrophils, macrophage, and non-immune cells (endothelial cells, fibroblasts), and enriched for type-1 immune (*CXCL9-11, IL6 IL1B, TNF*) and type-1 IFNs (*IFNB1, IFNA1, IFNA2, IFNA16)* signaling – the latter implicated in TB pathogenesis and having been previously demonstrated to contribute to neutrophil extracellular trap (NET) formation and subsequent *Mtb* proliferation (**Figure 7A,B** and **S5A-E**; Moreira-Teixeira et al., 2020).

**Figure 7.**
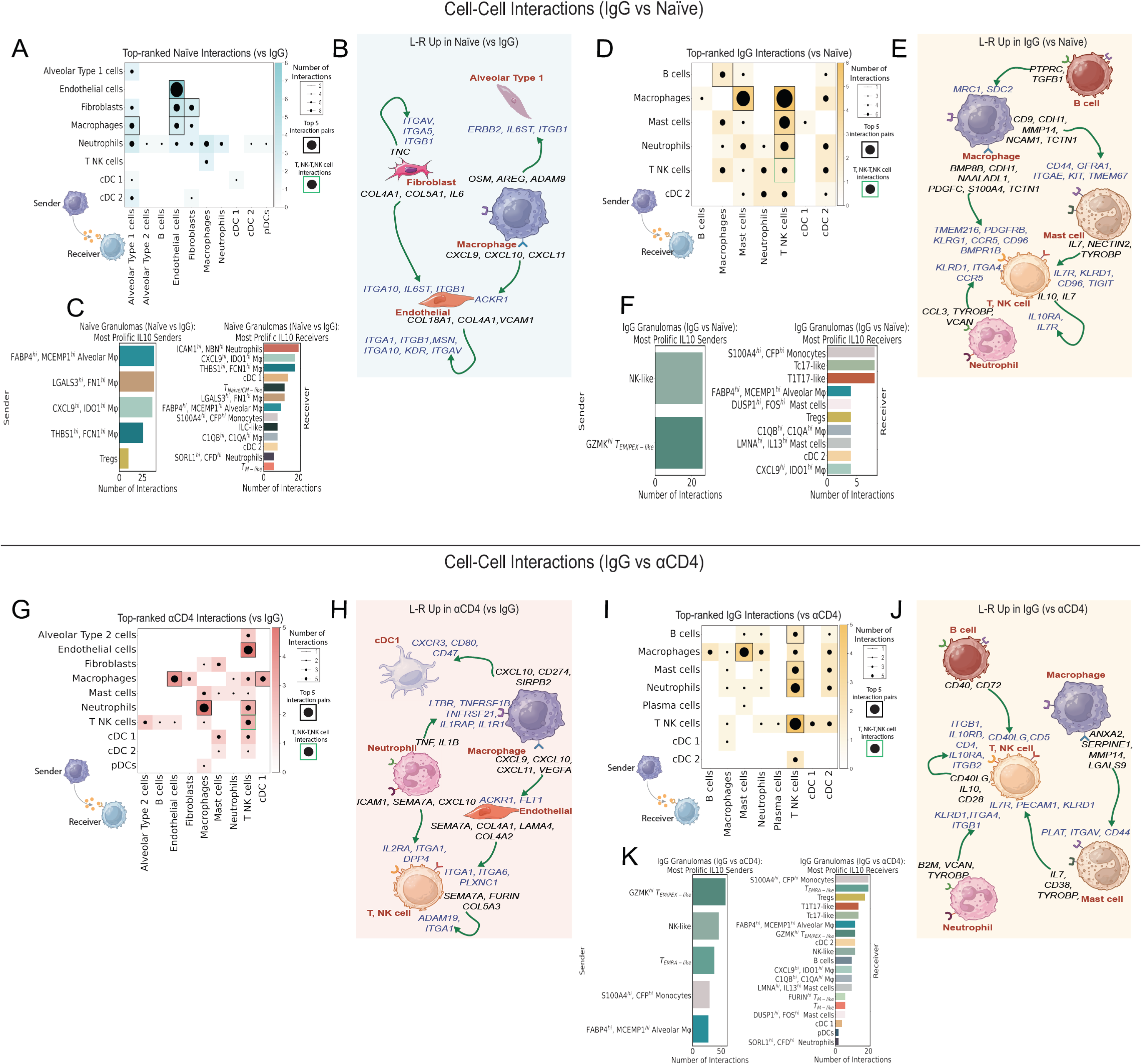
Differential cell-cell interactions in immunologically primed granulomas. **(A) (A)** Heatmap depiction of differential (naïve vs IgG) cell-cell interaction pairs among coarse cell types. Columns represent cell-cell interactions from the top-prioritized links – “sender” ligands and receptors differential L-R pairs specific to IgG or naïve granulomas. Heat map and dot size represent L-R interactions from the 50 top-prioritized links. Black rectangles indicate the top 5 interactions, based on number of interactions between two cell types, per NHP group. Green rectangles depict putative T, NK-T, NK interactions. **(B)** Cartoon depiction of (A) with differential L-R (i.e., top-prioritized linkages) specific to naïve granulomas. **(C)** Barplot depiction of differential cell-cell interactions among naïve granulomas. Left barplot depicts *IL10^+^*sender cellular subpopulations; right barplot represents *IL10RA/RB^+^*cell subpopulations. Receptor-ligand and inferred interaction pairs are derived from the top 200 top-prioritized linkages. **(D)** Similar heatmap to that of (A), highlighting linkages specific to IgG (vs naïve) granulomas. **(E)** Schematic representation of the differential L-R pairs unique to IgG granulomas from (D). **(F)** Barplot representation of differential *IL10-IL10RA/RB* interactions among IgG lesions, similar to that of (C). **(G)** Heatmap of αCD4 (vs IgG) granulomas**. (H)** Schematic representation of the differential L-R pairs unique to αCD4 granulomas from (G). **(I)** Heatmap of IgG (vs αCD4) granulomas. **(J)** Schematic representation of the differential L-R pairs unique to IgG granulomas from (I). **(K)** Barplot representation of differential *IL10-IL10RA/RB* interactions among IgG lesions, similar to that of (C).

To identify subpopulation-specific drivers of granuloma immune tone and cytokine flux, we quantified differential cell-cell interactions among all immune cell subpopulations (**Figure S5F,G**). This analysis identified several monocyte-derived subpopulations (*S100A4*^hi^, *CFP*^hi^ monocytes, *FABP4*^hi^, *MCEMP*^hi^ alveolar Mφ, *CXCL9*^hi^ *IDO1*^hi^ Mφ) “sending” the type-1 immune molecules *CXCL9-10*, and neutrophils (*ICAM1*^hi^, *NBN*^hi^ neutrophils, *SORL1*^hi^, *CFD^hi^* neutrophils) “sending” *TNF, CSF1,* and *CXCL10,* which targeted both innate and adaptive (e.g., *CXCR3^+^* T1T17-like) cellular subpopulations (**Figure S5F**). In addition to the upregulated type-1 immune factors, our subpopulation-specific cell-cell interaction analysis identified prominent monocyte-derived (*THBS1* ^hi^*, FCN1* ^hi^ Mφ; *LGALS3*^hi^, *FN1*^hi^ Mφ; *FABP4*^hi^, *MCEMP*^hi^ alveolar Mφ; *CXCL9*^hi^ *IDO1*^hi^ Mφ) *IL10* “senders” which targeted *ICAM1*^hi^, *NBN*^hi^ neutrophils, and several monocyte-derived subpopulations as “receivers” (**Figure 7C** and **S5F,G**). Collectively, these data may suggest type-1 immune signaling networks may promote the recruitment of adaptive immune cells and induce a pro-inflammatory immune response to mount an early anti-microbial response, whereas *IL10^+^* monocyte-derived subpopulations attempt to mitigate this inflammatory response via self-reinforcing innate-innate immune cell circuits.

In contrast to naïve lesions, our analysis of coarse-level cell types in IgG granulomas revealed T, NK cells as the dominant “receiver” cell type, and macrophages as the most prolific “senders,” communicating not only with T, NK cells, but also, strikingly, with mast cells (**Figure 7C,D** and **S5B,D,E)**. Outgoing macrophage-derived signaling was dominated by negative regulators of inflammatory response (*CD9, CD52, CDH1*), wound healing (*MMP14, S100A4*), and pro-angiogenic (*PDGFC*) signaling (Ackerman et al., 2019; Atkinson et al., 2007; Rashidi et al., 2018; Suzuki et al., 2009; **Figure S5B,E**). Outgoing mast cell signaling *IL7, TYROBP,*and *NECTIN2* largely targeted T, NK cell receivers expressing *TIGIT, IL7R, PECAM1, KLRD1,* suggesting immunoregulatory (*NECTIN2-TIGIT*) and homeostatic (*IL7-IL7R*) signaling axes among these cells. Our analysis also identified prominent B cell-macrophage communication via *TGFB1-SDC2* and *PTPRC-MRC1*, potentially suggesting that B cells contribute to macrophage polarization in the granuloma (Gong et al., 2012). Our subpopulation-specific cell-cell interaction analyses identified: (i) several monocyte-derived and mast cell subpopulations contributed to these wound-healing and anti-inflammatory signaling pathways and (ii) blunted type-1 immune network topologies (**Figure S5F,G**). There was also T, NK signaling to T,NK cells; these circuits were characterized by *IL10-IL10RA* and *IL7-IL7R*, suggesting that reinfection granulomas and the associated cytokine milieu and cellular composition promote self-reinforcing immunosuppressive and homeostatic regulatory T cell interactions (**Figure S7D-E;** Sun et al., 2011). A subpopulation-specific query of these cells revealed that GZMK^hi^ T_EM/PEX-like_ were one of the putative *IL10* “sender” populations and primarily targeted *IL10RA^+^ S100A4*^hi^, *CFP*^hi^ monocytes, Tc17-like, and T1T17-like cells. This analysis further defined a shift away from *IL10^+^*innate “sender” cellular subsets toward *IL10^+^* adaptive immune cell subpopulations: GZMK^hi^ T_EM/PEX-like_ and NK-like cells, relative to naive granulomas, with these two T, NK cell subsets targeting several T, NK cells subsets (Tc17-like, T1T17-like, Tregs) (**Figure 7F**).

In the absence of CD4^+^ T cells, TB granulomas were again dominated by pro-inflammatory neutrophil-derived and type-1 IFN signaling (**Figure 7G,H** and **S5H**, **S5J**-**L**). Outgoing neutrophil-derived signaling was enriched for type-1 immune signaling (*TNF, IL1B, CXCL10*), targeting macrophages and T, NK cells. There was a relative loss of mast cell signaling, including *IL7* signaling to *IL7R* expressing T, NK cells, suggesting a loss of homeostatic cycling of memory T, NK cell subsets, compared to IgG (**Figure 7I,J** and **S5I**). CD4-depletion lesions were further characterized by a relative loss of T, NK-T, NK signaling circuits involving “sender” ligands (*IL7, IL10, CD40LG, CD28*) and corresponding “receiver” receptors (*CD4, IL10RA, IL10RB, B, IL7R, ITGB1-2*) on T, NK cells (**Figure 7H,I** and **S5K-L**). Given the enrichment of *IL10* among coarse-level T, NK cells, we systematically queried all immune cell subpopulations to identify the putative *IL10* sender subpopulation(s), which revealed two terminally differentiated (GZMK^hi^ T_EM/PEX-like_, T_EMRA-like_) and one innate-like (NK-like) T cell subpopulation, as well as two monocyte-derived subpopulations (*S100A4*^hi^, *CFP*^hi^ monocytes, *FABP4*^hi^, *MCEMP*^hi^ alveolar Mφ) which “targeted” nineteen immune cell subpopulations, including eight T, NK cell subpopulations, suggesting the presence of CD4^+^ T cells is necessary for *CD8* T, NK cell immunomodulation and regulation in the TB granuloma (**Figure 7K** and **S5M,N**). Strikingly, IgG granulomas also demonstrated robust B cell signaling compared to αCD4 lesions, which lacked B cell contributions, among the top 50 prioritized linkages, to the granuloma cell-cell interactome (**Figure 7H,I**). Compared to αCD4 granulomas, IgG B cell “sender” ligands (*B2M, CD40, CD72, RPS19, TGFB1*) targeted four cell types, with two (*CD40, CD72*) of the five top-ranked ligands targeting T, NK cell receptors (*CD40LG, CD5*) – a potential consequence (direct or indirect) of CD4^+^ T cell depletion (Crotty, 2011; Sangster et al., 2003).

In summary, our systematic examination of the TB granuloma microenvironment following *Mtb* reinfection delineated distinct cellular circuitries, presenting a spectrum of responses – from amplification to dampening of the host inflammatory response – and underscoring the intricate balance in immune regulation associated with enhanced TB control.

## DISCUSSION

The TB granuloma represents a perturbed immunological niche where tissue resident and nascently recruited cells work together against microenvironmental stressors (e.g., *Mtb*, cellular enrichment/depletion, fibrosis, necrosis, inflammation, hypoxia) in an attempt to restore homeostasis. These responses can either promote bacterial control or dissemination, as well as tissue damage or preservation (Pagán and Ramakrishnan, 2018, 2015). In the present study, we leveraged PET-CT, microbiological assays, flow cytometry, and scRNA-seq of *Mtb* infected cynomolgus granulomas following primary *Mtb* infection and reinfection to identify the cellular features of protection that primary *Mtb* infection provides against *Mtb* reinfection and to examine how CD4^+^ T cell depletion before reinfection modulates host immunity. Collectively, our multi-modal dataset reveals global shifts in cellular composition, gene programming, and *Mtb* dynamics in primary *Mtb* infection and reinfection and nominates mechanisms by which CD4^+^ T cells contribute to a restrictive immunological niche. In doing so, our study yields new insights into the cellular, molecular, and niche features that support anti-*Mtb* activity or promote maladaptive immunity following natural infection – most critically, that CD4^+^ T cells act as homeostatic regulators of inflammation. It also identifies tissue-level cellular response mechanisms that can be targeted in future investigations for the development of improved prophylactics and cures.

Our high-dimensional examination of TB reinfection granulomas revealed underlying mechanisms governing granuloma cellularity and cytokine flux, as well as putative cell-cell interactions, providing insights into their roles in modulating anti-mycobacterial immunity. Illustratively, IgG lesions featured robust upregulation of immuno-modulatory and -regulatory genes (*IL10, PDCD1, TOX, TIGIT*) among lymphocyte-derived transcriptomes relative to primary infection (naïve) granulomas, which appeared to be CD4-dependent. Strikingly, cytokines canonically associated with protective TB immunity (*TNF*, *IFNG*) did not distinguish IgG granulomas from naïve or αCD4 lesions. These data corroborate our previous findings demonstrating that reinfection macaques concurrently infected with *Mtb* experienced significantly increased IL-10 secretion and relatively lower levels of TNF and IFNγ production (Cadena et al., 2018). Moreover, recent work assessing the efficacy of pulmonary mucosal BCG delivery identified IL-10^+^ T cells as the most robust correlate of protection (Dijkman et al., 2019). In both studies (Cadena et al., 2018; Dijkman et al., 2019), the source of IL-10 production among T cells was unknown. Our present work expands these findings, identifying several T, NK cell subpopulations, including terminally differentiated and cytotoxic *CD8-*enriched subpopulations as putative sources of *IL10* production in the TB granuloma. Altogether our data demonstrate a shift in reinfection granuloma cytokine flux, cellularity, and programming, with T, NK cells biasing towards *CD8-*enriched immunoregulatory phenotypes.

Our cell-cell interaction analyses further identified roles for the immunoregulatory molecules *IL10* and *TIGIT* – expressed among T, NK cells – following *Mtb* reinfection. Significantly, these molecules were absent among naïve and CD4-depleted signaling networks, suggesting that following immune priming, *CD8*^+^ T, NK cells require CD4^+^ T cell help (direct or indirect) to engage in self- and non-self-immunoregulatory circuits. Critically, the immunoregulatory molecule PD-1 has been demonstrated to promote host-resistance to TB, with checkpoint inhibitors (PD-1 blockade) exacerbating TB disease and immunopathology via overproduction of IFNγ from CD4^+^ CXCR3^+^, KLRG1^-^, CX3CR1^-^ T cells and elevated infiltrates of pro-inflammatory CD8^+^ T cells (Kauffman et al., 2021; Sakai et al., 2016); this suggests that the immunoregulatory circuits we identified among reinfection T, NK cells may be attempting to balance granuloma equilibria and mitigate tissue damage. While PD-1, IL-10, TIGIT, and other immunosuppressive molecules may mitigate inflammatory pathophysiology associated with TB, they may also inadvertently foster an environment conducive to *Mtb* persistence (Redford et al., 2011, 2010; Wong et al., 2020); this highlights the importance of balanced interplay between IL-10, among other immunoregulatory molecules (e.g., TIGIT, PD-1), and type-1 immunity in anti-TB immunity (Gideon et al., 2015).

Our analyses revealed that the immunoregulatory molecules *TIGIT, IL10,* and *PDCD1* were upregulated in the *CD8^+^* GZMK^hi^ T_EM/PEX-like_ T, NK cell subpopulation. A comparison of GZMK^hi^ T_EM/PEX-like_ frequencies revealed that immunologically primed (IgG) animals experience significantly elevated recruitment (or retention) of GZMK^hi^ T_EM/PEX-like_ cells relative to naïve or αCD4 lesions, suggesting that GZMK^hi^ T_EM/PEX-like_ localization is CD4^+^ T cell-dependent. Although GZMK^hi^ T_EM/PEX-like_ cells were enriched in granuloma with dampened type-1 immune cellularity and inflammatory response, scRNA-seq analyses of disparate pathologies (e.g., rheumatoid arthritis, cancer, ulcerative colitis, viral infection) and tissues have suggested GZMK^+^ CD8^+^ T cells promote and potentiate inflammatory sequelae (Jonsson et al., 2022; Thomas et al., 2021; Zheng et al., 2021). Moreover, these scRNA-seq studies have demonstrated GZMK^+^ CD8^+^ TCRs are highly clonal and restricted to sites of inflammation, potentially suggesting that these cells differentiate at the site of disease or become differentiated before recruitment (Cai et al., 2022; Jonsson et al., 2022). An analysis of GZMK^+^ CD8^+^ transcriptomes and TCRs derived from TB pleural fluid showed that clonally expanded GZMK^+^ CD8^+^ cells were restricted to pleural fluid and absent in peripheral blood (Cai et al., 2022). Furthermore, *in vitro* experiments leveraging *Mtb*-infected macrophages demonstrated purified GZMK has both cytotoxic and anti-microbial effector functions. While the absence of TCR and PBMC sampling in our study precluded us from identifying the origin of GZMK^+^ CD8^+^ T cells, granuloma GZMK^hi^ T_EM/PEX-like_ cells had only low expression of T cell migration factors (CCR7, *SELL*, *CXCR5*), and upregulated expression of genes canonically associated with chronic inflammation and immunoregulation potentially suggesting: (i) granuloma GZMK^hi^ T_EM/PEX-like_ cells have intra-compartment/lesion migratory potential, and (ii) GZMK^hi^ T_EM/PEX-like_ cells acquire a terminally differentiated phenotype at the site of infection. Collectively, these data suggest that GZMK^hi^ T_EM/PEX-like_ recruitment (or retention), differentiation, and state may be CD4^+^ T cell-dependent, further supporting a critical role for CD4^+^ T cells in balancing pro- and anti-inflammatory immunity.

Upon *Mtb* exposure, myeloid-derived cells, namely alveolar macrophages, sense and phagocytose invading bacilli, activating cell-intrinsic anti-microbial and inflammatory pathways, thus deviating from homeostasis to clear infection (Cohen et al., 2018; Ravesloot-Chávez et al., 2021). Following failure to clear initial infection, bacilli disseminate to lymphoid tissues and persist in non-restrictive cellular niches (Ganchua et al., 2018). Indeed, in the absence of Th1 activation of myeloid-derived cells, microbial growth remains unrestricted; thus, the ‘central dogma’ of anti-mycobacterial immunity has placed significant weight on IFN-γ and TNF Th1 CD4^+^ T cell-mediated immunity (Nunes-Alves et al., 2014). In addition to CD4^+^ IFN-γ and TNF production, T cells secrete the immunomodulatory cytokine IL-10 which dampens both adaptive and innate immunity, downregulating MHCs, reactive oxygen and nitrogen intermediates, and type-1 chemokines (e.g., CXCL10), as well as inhibiting phagosome maturation among innate cells (Kessler et al., 2017; O’Leary et al., 2011; Rene de Waal Malefy et al., 1991). In keeping with these findings, we observe reinfection (IgG) granulomas, enriched for *IL10* expressing T, NK cells, experienced significant blunting of type-1 inflammation and type-1 IFN signaling relative to primary infection (naïve) and αCD4 granulomas. Furthermore, our findings demonstrate IgG monocyte-derived transcriptomes downregulate type-1 chemokines (e.g., CXCL9-11), as well as the IFN-γ and TNF response pathways – potentially a nuanced mechanism wherein the host attempts to achieve an equilibrium between protective immunity and tissue preservation (Gazzinelli et al., 1996; Sun et al., 2009). This modulation, while potentially mitigating tissue damage, may inadvertently reduce the host’s capacity to sterilize phagocytosed *Mtb*. Notably, however, our *Mtb* barcode data demonstrate reinfection (IgG) animals had significantly fewer tissues sharing *Mtb* barcodes, indicative of reduced bacterial dissemination compared to naïve granulomas, and that in the absence of CD4^+^ T cells, reinfection macaques had significantly elevated sharing of bacterial barcodes, highlighting the pivotal role of CD4^+^ T cells in modulating an effective host response that can mitigate bacterial establishment during reinfection and *Mtb* dissemination between compartments. The intricate host-pathogen dynamics in TB, as reflected by these findings, necessitate a comprehensive understanding not only of the evident immune markers but also of the broader roles that macrophages play within the inflammatory landscape.

Macrophage sensing and phagocytosis of *Mtb* during acute infection triggers the production of pro-inflammatory chemokines (CXCL1, CXCL2) and cytokines that promote vascular permeability, upregulation of adhesion molecules, and subsequent neutrophil recruitment (Cai et al., 2010; Phillipson and Kubes, 2011). Our scRNA-seq analyses uncovered previously unappreciated neutrophil heterogeneity – cell types that have been underrepresented in droplet-based single-cell profiling of TB (Esaulova et al., 2020; Pisu et al., 2021) – in the TB granuloma, including the identification of two neutrophil subsets (*ICAM1*^hi^, *NBN*^hi^ neutrophils; *SORL1*^hi^, *CFD*^hi^ neutrophils), with differential pathway activation and phenotypic signatures. Notably, our data demonstrate that hypoxia- and inflammatory-enriched naïve and αCD4 granulomas have significant neutrophilia, and significant induction of an IFN-responsive module associated with active TB, which may contribute to tissue inflammation and lung structural damage and the formation of an *Mtb*-permissive niche (such as caseum), thus promoting *Mtb* growth and dissemination (Berry et al., 2010; Lovewell et al., 2021). Critically, our data demonstrate reinfection (IgG) granulomas – characterized by reduced neutrophilia – do not support the same level of bacterial growth or dissemination as naïve lesions. They also show that inhibition of *Mtb* outgrowth and dissemination post-reinfection is at least partially CD4^+^ T cell-dependent and independent of CD4^+^ T cell-mediated induction of myeloid IFNγ- and TNF-response pathways.

In line with our findings, which demonstrated that 4-week post-primary infection (i.e., naïve NHP cohort) granulomas feature robust type-1 immune induction (e.g., *IL1B, CXCL9-11*) and signaling, previous research has shown that the chemokines CXCL9-11 are enriched among 4 week primary granulomas and that those granulomas feature elevated CXCR3^+^ T cell frequencies – putative sources of IFNγ and TNF (Gideon et al., 2022; Lin et al., 2006; Mehra et al., 2010). These findings highlight the critical role of CXCL9-11 during early *Mtb* infection (before immune priming), where they promote the recruitment of protective lymphocyte populations, such as CXCR3^+^ T cells; however, their overexpression may drive TB sequelae via self-reinforcing pro-inflammatory myeloid (neutrophil, macrophage)-T, NK cell (CXCR3^+^ IFN- γ^+^ TNF^+^ T1T17-like) circuits, thus potentiating pro-inflammatory response mechanisms and subsequent bacterial dissemination (Esaulova et al., 2020; Sakai et al., 2016). Indeed, the chemokines CXCL9-11 have been identified as potential biomarkers of TB severity, with overexpression reported in human study participants with active TB; however, the temporal dynamics of CXCL9-11 expression during active human TB are unclear as time of infection is frequently unknown (Bhattacharyya et al., 2018; Kumar et al., 2021; Lee et al., 2015; Nonghanphithak et al., 2017; Ruhwald et al., 2007). While there is substantial evidence suggesting that excessive CXCL9-11 may contribute to the early anti-*Mtb* immunity or immunopathology, other studies suggest that CXCL9-11 could serve as markers of trained immunity following BCG vaccination, potentially acting as an initial barrier against invading bacilli, as well as correlate of protection (Joosten et al., 2018). Nevertheless, the precise role of CXCL9-11 in immunity requires further exploration. Future work leveraging single-cell multi-omics data (e.g., scATAC-seq) in BCG-vaccinated and *Mtb*-infected macaques could help determine whether, and possibly when, elevated type-1 immune signaling is indicative of innate training or a correlate of TB pathophysiology and chronic inflammatory stimuli.

In summary, we provide the first in-depth characterization of primary infection, reinfection, and CD4^+^ T cell-depleted reinfection macaque granulomas, identifying potential mechanisms by which CD4^+^ T cells contribute to anti-mycobacterial immunity. Our high-dimensional characterization of granuloma transcriptomes and multimodal analyses reveal cellular networks in which CD4^+^ T cells regulate pro- and anti-inflammatory gene programming and cell-cell signaling networks to limit inflammatory sequelae, as well as bacterial establishment, growth, and dissemination. These findings expand beyond the limited purview of the TB ‘central dogma,’ demonstrating that CD4^+^ T cells act not only as effectors secreting IFN-γ and TNF but also as homeostatic regulators, orchestrating both pro- and anti-inflammatory immunity, thus leading to a more nuanced understanding of protective immunity against TB disease, and broader understanding of how CD4^+^ T cells modulate the immune response in reinfection events.

## Limitations of the study

While our study provides pivotal insights into the role of CD4^+^ T cells in the context of *Mtb* reinfection, it is important to acknowledge several inherent limitations associated with our design and experimental power, including: (i) the need to use a 4–5-month drug treatment to clear primary infection; and, (ii) limited number of NHPs and granulomas for scRNA-seq. Additionally, our study design does not enable us to compare the naïve and αCD4 groups directly since we did not have sufficient numbers of animals or granulomas and cannot disentangle shifts due to immunological priming from those due to CD4^+^ T cell depletion.

## SUPPLEMENTAL INFORMATION

Supplemental information can be found online.

## ACKNOWLEDGEMENTS

We extend our appreciation to the technical and veterinary teams of the Flynn, Shalek, and Fortune laboratories for their dedication and assistance. We are particularly indebted to the University of Pittsburgh’s Division of Laboratory Animal Research and the Unified Flow Core for their animal husbandry and support. Moreover, we are grateful to the NIH Nonhuman Primate Reagent Resource for providing essential antibody reagents (the CD4 depleting mAb used in this study was developed and provided by the NIH Nonhuman Primate Reagent Resource (ORIP P40 OD028116, U24 AI126683) and their expert guidance on their application. This work was supported in part by the Bill and Melinda Gates Foundation (A.K.S.), NIH R01 AI114674 (Flynn and Fortune), and by NIH contract IMPAc-TB; 75N93019C00071 (Fortune, Flynn, Shalek).

## AUTHOR CONTRIBUTIONS

Conceptualization, J.L.F, S.M.F., and A.K.S.; Data curation J.D.B., S.K.C.G., M.C., H.P.G., R.F.-O.; formal analysis J.D.B., S.K.C.G., S.K.N., P.M., M.C., D.W.; Investigation J.D.B., S.K.C.G., S.K.N., P.M., M.C., D.M., S.N., J.R., H.P.G.; Visualization J.D.B., P.M., M.C.; Funding acquisition J.L.F, S.M.F., and A.K.S; Supervision J.L.F, S.M.F., A.K.S., and B.B.; Writing – original draft J.D.B., D.M., J.L.F, S.M.F., and A.K.S.; Writing – review & editing J.D.B., S.K.C.G., S.K.N., M.C., D.M., S.N., J.R., J.L.F, S.M.F., and A.K.S.

## DECLARATIONS OF INTERESTS

A.K.S. reports compensation for consulting and/or scientific advisory board membership from Honeycomb Biotechnologies, Cellarity, Ochre Bio, Relation Therapeutics, FL86, IntrECate Biotherapeutics, Bio-Rad Laboratories, Senda Biosciences and Dahlia Biosciences unrelated to this work. S.M.F. reports compensation for board of directors membership from Oxford Nanopore unrelated to this work.

## KEY RESOURCES TABLE

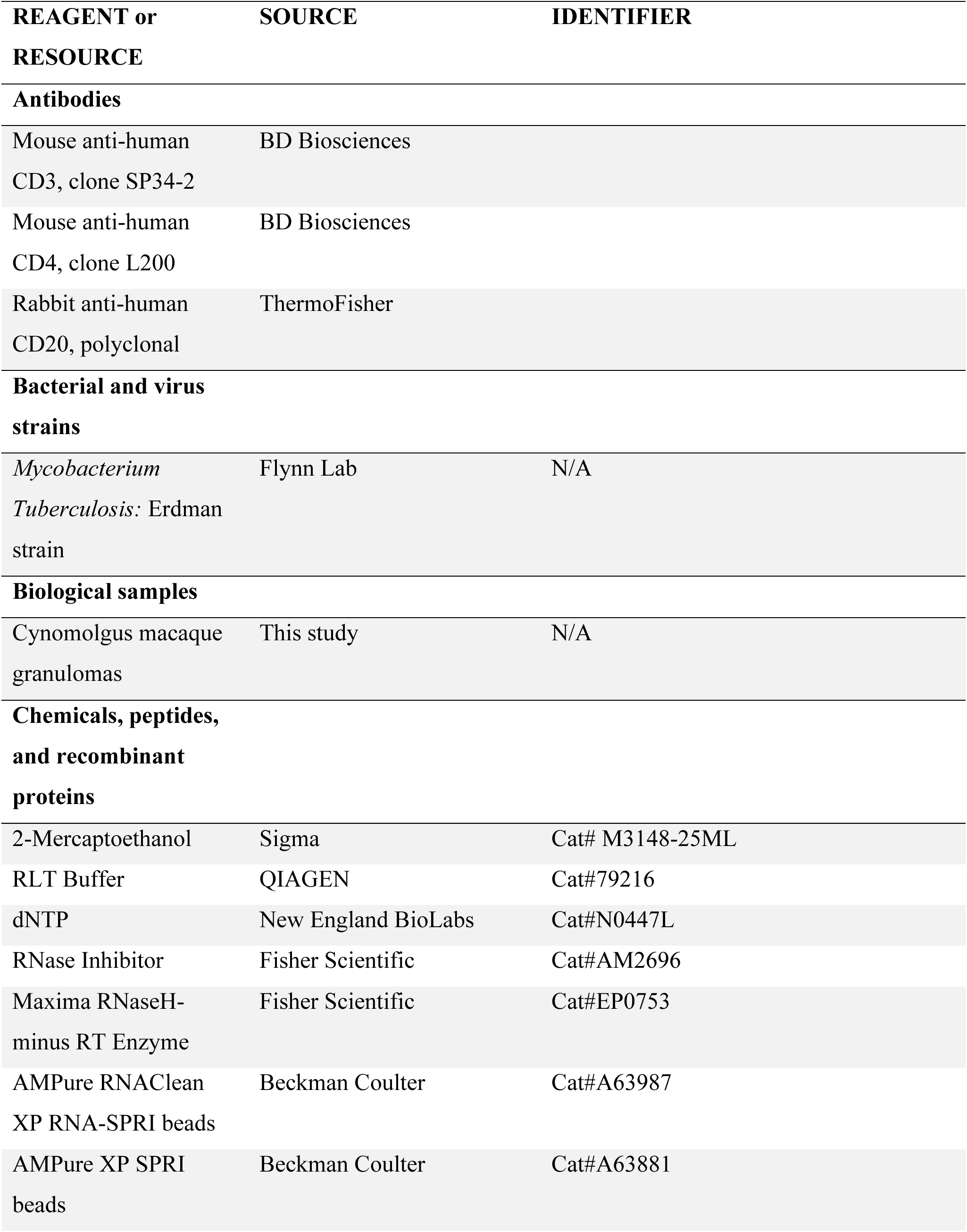
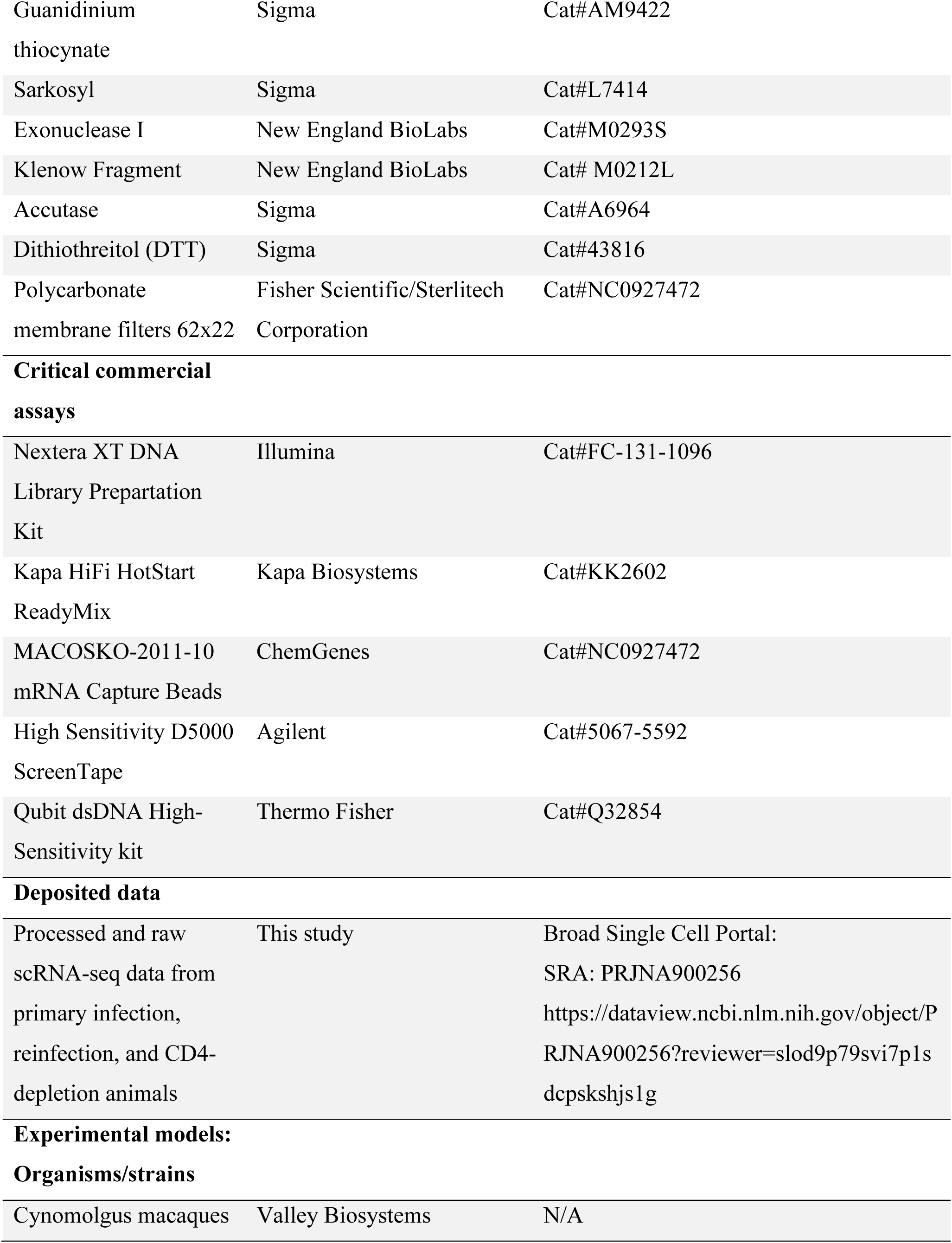
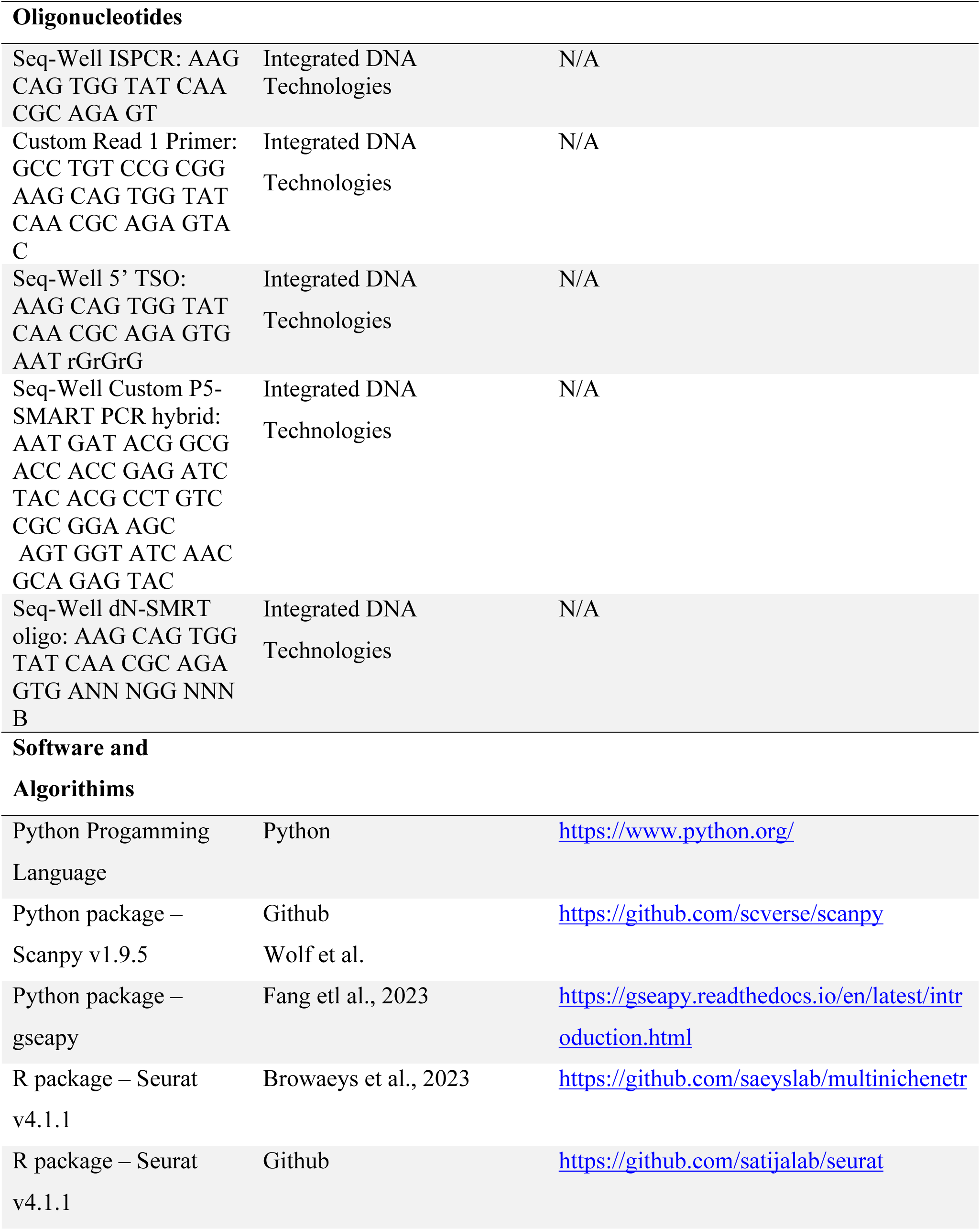
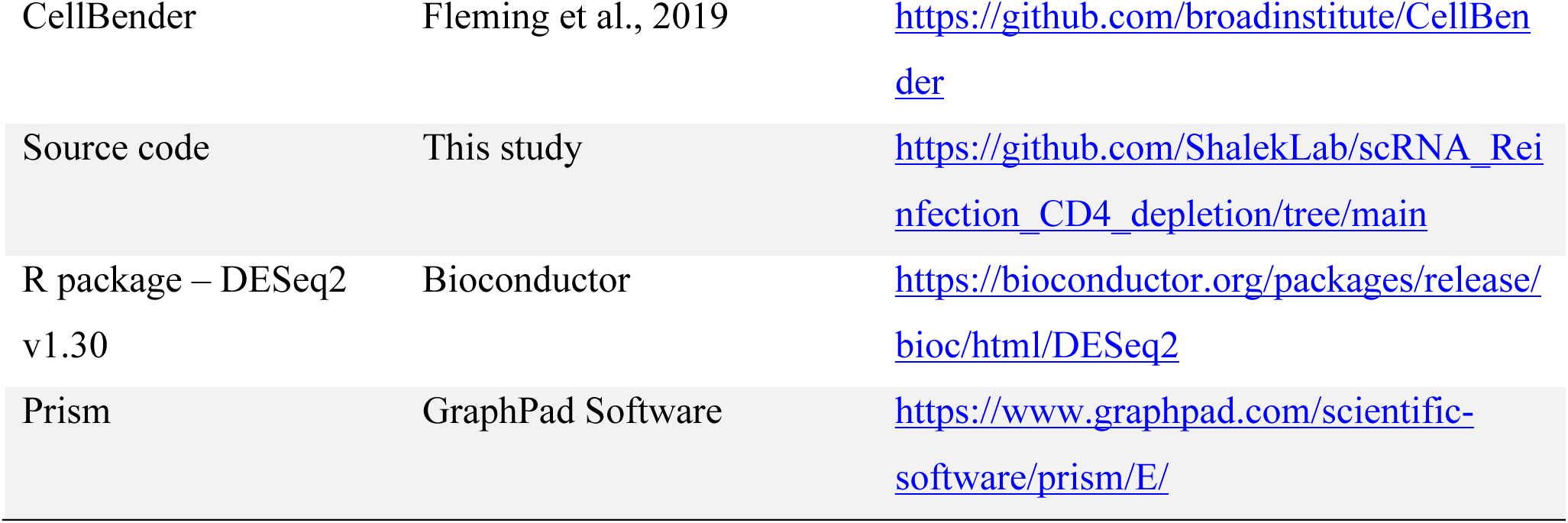

## RESOURCE AVAILABILITY

Further information and requests for resources and reagents should be directed to and will be fulfilled by the lead contact Dr. JoAnne L Flynn.

### Data and Code Availability

scRNA-seq data is publicly available for download and visualization via the Alexandria and Broad Institute Single Cell Portal upon publication. Accession numbers and links are listed in the key resources table. All original code has been deposited at GitHub (https://github.com/ShalekLab) and is publicly available as of the date of publication. Any additional information required to reanalyze the data from this study is available from the lead contact upon request. All the data generated in support of the reported findings can be found at: https://fairdomhub.org/studies/1239 by the time of publication, including scRNASeq, barcoding, flow cytometry and PET-CT imaging data

## EXPERIMENTAL MODEL AND SUBJECT DETAILS

### Research Animals

All experimental manipulations, protocols, and care of the animals were approved by the University of Pittsburgh School of Medicine Institutional Animal Care and Use Committee (IACUC). The protocol assurance number for our IACUC is A3187-01. Our specific protocol approval number for this project is 15066174. The IACUC adheres to national guidelines established in the Animal Welfare Act (7 U.S.C. Sections 2131–2159) and the Guide for the Care and Use of Laboratory Animals (8^th^ Edition) as mandated by the U.S. Public Health Service Policy.

All macaques used in this study were housed at the University of Pittsburgh in rooms with autonomously controlled temperature, humidity, and lighting. Animals were singly housed in caging at least 2 square meters apart that allowed visual and tactile contact with neighboring conspecifics. The macaques were fed twice daily with biscuits formulated for nonhuman primates, supplemented at least 4 days/week with large pieces of fresh fruits or vegetables. Animals had access to water *ad libitem*. Because our macaques were singly housed due to the infectious nature of these studies, an enhanced enrichment plan was designed and overseen by our nonhuman primate enrichment specialist. This plan has 3 components. First, species-specific behaviors are encouraged. All animals have access to toys and other manipulata, some of which will be filled with food treats (e.g., frozen fruit, peanut butter, etc.). These are rotated on a regular basis. Puzzle feeders foraging boards, and cardboard tubes containing small food items also are placed in the cage to stimulate foraging behaviors. Adjustable mirrors accessible to the animals stimulate interaction between animals. Second, routine interaction between humans and macaques are encouraged. These interactions occur daily and consist mainly of small food objects offered as enrichment and adhere to established safety protocols. Animal caretakers are encouraged to interact with the animals (by talking or with facial expressions) while performing tasks in the housing area. Routine procedures (e.g., feeding, cage cleaning, etc) are done on a strict schedule to allow the animals to acclimate to a routine daily schedule. Third, all macaques are provided with a variety of visual and auditory stimulation. Housing areas contain either radios or TV/video equipment that play cartoons or other formats designed for children for at least 3 hours each day. The videos and radios are rotated between animal rooms so that the same enrichment is not played repetitively for the same group of animals.

All animals are checked at least twice daily to assess appetite, attitude, activity level, hydration status, etc. Following *M. tuberculosis* infection, the animals are monitored closely for evidence of disease (e.g., anorexia, weight loss, tachypnea, dyspnea, coughing). Physical exams, including weights, are performed on a regular basis. Animals are sedated prior to all veterinary procedures (e.g., blood draws, etc.) using ketamine or other approved drugs. Regular PET CT imaging is conducted on most of our macaques following infection and has proved very useful for monitoring disease progression. Our veterinary technicians monitor animals especially closely for any signs of pain or distress. If any are noted, appropriate supportive care (e.g. dietary supplementation, rehydration) and clinical treatments (analgesics) are given. Any animal considered to have advanced disease or intractable pain or distress from any cause is sedated with ketamine and then humanely euthanatized using sodium pentobarbital.

## METHOD DETAILS

### Necropsy Procedures

Procedures done during necropsy have been previously described (Lin et al., 2009, 2006). Briefly, 1-3 days prior to necropsy, a PET CT scan was taken and used to identify the location and metabolic activity (FDG avidity) of granulomas and lymph nodes; this scan was used as a map to aid in the individual identification and excision of these samples during necropsy. On the day of necropsy, macaques were humanely sacrificed with sodium pentobarbital and terminally bled. Individual granulomas, thoracic and peripheral lymph nodes, lung tissue, spleen and liver were all excised and homogenized separately into single cell suspensions. New granulomas determined by PET-CT and uninvolved lung lobes (no granuloma present in the lobe) were enzymatically homogenized using a human tumor dissociation kit (Miltenyi Biotec) and a gentleMACS Dissociator (MiltenyiBiotec) following manufacturer’s protocols. Homogenates were aliquoted for plating on 7H11 agar for bacterial burden, freezing for DNA extraction and staining for flow cytometry analysis. Any remaining samples were frozen for future use. Homogenates were plated in serial dilutions on 7H11 medium and incubated at 37°C/5% CO_2_ for 3 weeks before enumeration of CFU.

### Isolation of genomic DNA from bacteria

DNA extraction was performed on granuloma and lymph node homogenates, as well as their scrapates (scraped colonies that grew on 7H11 agar plates) for library identification as described previously (Lin et al., 2014). Briefly, a small aliquot of the homogenate or scrapate were vortexed with 0.1mm zirconia-silica beads (BioSpec Products, Inc.) and subsequently extracted twice with phenol chloroform isoamyl alcohol (25:24:1, Sigma-Aldrich) before precipitating DNA with molecular grade 100% isopropanol (Sigma-Aldrich) and 3M sodium acetate (Sigma-Aldrich) and resuspending in nuclease-free water (Invitrogen).

### Library identification

Identification of library DNA tags have been previously described (Cadena et al., 2018). Briefly, DNA was amplified by PCR for 24-36 cycles before using in the NanoString nCounter assay (NanoString Technologies) with custom designed probes (Martin et al., 2017). New granulomas after reinfection were identified by PET-CT scan comparing pre- and post-reinfection scans and verified by presence of Library S barcodes.

### Intracellular cytokine staining and flow cytometry

Because of the abundance of *Mtb* antigens already present in granulomas and involved lymph nodes (Gideon et al., 2015), these samples were not further stimulated with *Mtb* peptides (Gideon et al., 2015). Due to low number of cells, uninvolved lung lobes were also not stimulated. All samples were incubated in the presence of Brefeldin A (GolgiPlug, BD Biosciences) at 37°C/5% CO_2_ for 3.5-4 hours prior to staining. Cells were stained with a viability marker (LIVE/DEAD fixable blue dead cell stain kit, Invitrogen) and surface and intracellular markers. Surface markers include CD3 (clone SP34-2, BD Pharmingen), CD4 (Clone L200, BD Horizon), CD8 (Clone SK1, BD Biosciences) and CD20 (Clone 2H7, eBioscience).

### Animals, infections, CD4 depletion and disease tracking by PET CT

Nineteen cynomolgus macaques (*Macaca fasicularis*) with age range of 5.8-9.1 years were obtained from Valley Biosystems (Sacramento, California). Animals were placed in quarantine for 1 month where they were monitored to ensure good physical health and no prior *Mtb* infection. All animals were infected with Library P DNA-tagged *Mtb* Erdman via bronchoscopic instillation as previously described (Capuano et al., 2003; Lin et al., 2009). Thirteen macaques received *Mtb* library P as the first infection. Granuloma formation, lung inflammation and overall disease was tracked using ^18^F-fluorodeoxyglucose (FDG) PET-CT every 4 weeks throughout infection. PET-CT scans were analyzed using OsiriX viewer as previously described with a detection limit of 1mm (White et al., 2017). The first infection was followed for 9 weeks before drug-treating all 13 macaques. Based on our previous study (Lin et al., 2012b), exacerbation of TB disease occurs after CD4^+^ T cell depletion, thus to facilitate identification of new granulomas arising from the second infection we opted to treat all macaques with anti-TB drugs. Macaques were given anti-TB drugs orally once daily for 4-5 months (RIF 20mg/kg; INH 15mg/kg; ETH 50mg/kg; PZA 150mg/kg) (Lin et al., 2012a). Compliance ranged from 97-100%. The 13 macaques were matched by PET CT for disease status and randomized into 2 cohorts: CD4^+^ T cell depletion (n=7) and IgG control (n=6). After resting for 4 weeks after drug treatment, CD4R1 (50mg/kg, NHP Reagent Resource), a rhesus recombinant CD4^+^ T cell-depleting antibody, was administered subcutaneously in 4 animals and intravenously in 3 animals 1 week before the second infection with *Mtb* Library S and then was administered intravenously every 2 weeks until necropsy. CD4^+^ T cell depletion was monitored by flow cytometry in the blood and complete blood count weekly. To measure CD4^+^ T cell depletion in tissues, a peripheral lymph node biopsy was performed before CD4^+^ T cell depletion and the CD4^+^ T cell level was compared with a peripheral lymph node from the same macaque obtained at necropsy. Macaques from the IgG control group received rhesus recombinant IgG1 control antibody (50mg/kg, NHP Reagent Resource) following the same timeline of the CD4^+^ T cell-depletion group. Six macaques were included as naïve controls infected with *Mtb* Library S only.

Macaques received 4-12 CFU of *Mtb* Library P for the first infection and 8-22 CFU of *Mtb* Library S for the second infection (or the first infection for the naïve monkeys). Dose was calculated from colony counts after plating an aliquot of the infection inoculum on 7H11 agar plates and incubating for 3 weeks at 37°C/5% CO_2_.

### Antibody validation

To test whether the anti-CD4^+^ depletion antibody masks CD4 receptors, peripheral blood mononuclear cells (PBMC) were incubated with 1X (0.77 mg/ml, the calculated concentration of αCD4 antibodies in blood of macaques given a dose of 50mg/kg), 0.25X and 4X concentration of CD4 T cell-depleting antibody for 30 minutes at 37°C before surface staining with CD3 (clone SP34-2, BD Pharmingen), CD4 (Clone L200, BD Horizon) and CD8 (Clone RPA-T8, BD Biosciences) surface markers. PBMCs that were not incubated with the αCD4 antibody were included as a control. Data was acquired using the LSR II (BD) and analyzed using FlowJo software v10.6.1 (BD).

### Single-cell RNA-sequencing (scRNA-seq) and alignment

Massively parallel scRNA-seq was performed using the Seq-Well S^3^ platform, as previously described (Gierahn et al., 2017; Hughes et al., 2020). Approximately 15,000-20,000 cells were loaded onto Seq-Well arrays equipped with uniquely barcoded mRNA capture beads (ChemGenes). Cells were allowed to settle by gravity into wells for 10 minutes, after which the arrays were washed with PBS and serum-free RPMI. Arrays were then sealed with a semi-permiable polycarbonate membrane and incubated at 37°C for 30 minutes. Cell lysis was achieved by incubating the sealed arrays in a lysis buffer (5 M guanidine thiocyanate, 10 mM EDTA, 0.1% BME, and 0.1% sarkosyl) for 20 minutes. Subsequently, the arrays were rocked in hybridization buffer (2M NaCl, 8% v/v PEG8000) for 40 minutes. After membrane removal, the arrays were washed with in Seq-Well wash buffer (2M NaCl, 3 mM MgCl2, 20 mM Tris-HCl, and 8% v/v PEG8000) to collect the mRNA capture beads. Reverse transcription was conducted at 52°C using Maxima H Minus Reverse Transcriptase (ThermoFisher), and excess primers were removed with an Exonuclease I digestion (New England Biolabs). Whole transcriptome amplification (WTA) was achieved via PCR using KAPA Hifi PCR Mastermix (Kapa Biosystems). The WTA product was purified using Agencourt Ampure beads (Beckman Coulter). For sequencing, dual-indexed 3’ DGE libraries were prepared using Nextera XT (Illumina) and sequenced to depth on the NovaseqS4 platform with a paired-end read structure (R1: 20 bases; I1: 8 bases; I2: 8 bases; R2: 50 bases) using custom sequencing primers. Transcript reads were tagged for cell barcode and UMI utilizing DropSeqTools v1.12 (Macosko et al., 2015). These tagged sequencing reads were subsequently aligned to the *Macaca fascicularis* v5 genome (https://useast.ensembl.org/Macaca_fascicularis/Info/Index) using the Dropseq-tools pipeline on the Terra platform (app.terra.bio). By collapsing the aligned reads based on barcode and UMI sequences, we derived digital gene expression matrices for each array, covering 10,000 barcodes.

## QUANTIFICATION AND STATISTICAL ANALYSIS

### Statistical analysis on macaque samples (depletion and CFU)

Test for normality was conducted with a Shapiro-Wilk test. For assessment of depletion over time, mixed effects model with Dunnett’s multiple comparison test adjusted for the comparison of IgG vs anti-CD4 and IgG vs naïve. For pre- and post-PET data, two-way ANOVA with Bonferroni multiple comparisons test. For paired analyses, Wilcoxon matched-pairs signed rank tests were used. For comparison of three groups (IgG vs naïve and IgG vs αCD4), either one-way ANOVA with Dunnett’s multiple comparisons or Kruskal-Wallis with Dunn’s multiple comparisons were used dependent on normality.

### Data processing and quality control

For individual granuloma, collapsed gene expression matrices containing 10,000 barcodes were subject to CellBender to estimate the fraction of ambient RNA contaminating cell transcriptomes. The CellBender “remove-background” function was then applied using default parameters. Individual CellBender “corrected” matrices were then subject to Scrublet, using default parameters, to identify putative doublets. Transcriptomes with a doublet_score >0.30 were removed from downstream analyses. Sample-specific gene expression matrices were then combined and analyzed using Scanpy (version 1.8.2). Transcriptomes were filtered using min_genes > 300, min_counts>500, mitochondrial_threshold=0.05, and genes expressed in fewer than 10 cells were removed. Gene expression counts were normalized using default Scanpy parameters (i.e., log_2_(TP10K+1)).

### Dimensionality reduction and batch correction

Following preliminary filtering processes, we performed coarse-level cell type clustering and iterative sub-clustering to annotate cell types and identify low-quality transcriptomes (e.g., doublets) not identified or removed during preliminary quality control processing, respectively. The top 2,000 highly-variable genes – identified using the Scanpy “highly_variable_genes function” – were used for dimensionality reduction. Following variable gene selection, these data were subject to scaling, principal component analysis (PCA), integration to mitigate sample-specific batch effects, and Leiden-based clustering. More specifically, data were scaled to 10, and the top 19 principal components (PCs) were used for dimensionality reduction. PCs were used to construct a neighborhood graph using the scanpy.pp.neighbors function, setting n_neighbors=40 and using the top 19 PCs. Leiden-based clustering was then implemented, setting the resolution= 1.0, which identified 26 distinct clusters.

### Cell clustering and annotation

From these 26 clusters we identified 14 coarse cell types. The Leiden resolution=1.0 failed to distinguish between several cell types (e.g., cDCs 1 and pDCs; alveolar type-1 and alveolar type 2 cells). As a result, these preliminary coarse-level cell types were not used as the final reference but instead used to inform sub-clustering. All coarse-level cell types (e.g., T, NK cells, macrophages) were subject to sub-clustering to remove low-quality cells. Transcriptomes classified as doublets featured elevated expression of genes derived from distinct cell ontologies. These doublets were excluded from downstream analyses.

Following quality control processing, our data set comprised 88,360 high-quality transcriptomes, which were annotated as 15 distinct cell types, including: alveolar type-1 cells, alveolar type 2 cells, B cells, ciliated cells, endothelial cells, eosinophils, fibroblasts, macrophages, mast cells, neutrophils, plasma cells, T, NK cells, cDCs 1, cDCs 2, and pDCs. Among cDCs 1, cDCs 2, pDCs, and plasma cells additionally subset diversity was not found, as such these coarse-level annotations are equivalent to cellular subsets. The major cell populations alveolar type-1 cells, alveolar type 2 cells, B cells, ciliated cells, endothelial cells, eosinophils, fibroblasts, macrophages, mast cells, neutrophils, and T, NK cells underwent further sub-clustering to discern cellular subtypes. Sub-clustering resolution was determined by selecting the most stable/robust silhouette score that uncovered biologically relevant and/or known cell subsets (e.g., Tregs).

### Differential abundance analysis of scRNA-seq cell type and subset frequencies

To identify differential cell type frequencies across naïve, IgG, and αCD4 granuloma, we implemented three distinct statistical frameworks: (1) scCODA, (2) the Mann-Whitney U-test, and (3) Fischer’s exact test.

One inherent challenge in scRNA-seq data is the compositional nature of cell proportions – they are not mutually exclusive. Illustratively, the elevation of one cell subset’s proportion inherently diminishes the proportions of others due to the requirement that all proportions sum to one (e.g., antibody-mediated CD4^+^ T cell depletion results in elevated frequencies of CD8A^+^ T cells among T, NK cells). To address these limitations, we implemented scCODA, a statistical framework rooted in a Bayesian hierarchical model, which is adept at dissecting cell type co-dependencies and the low inputs typically associated with scRNA-seq data, thus ensuring that observed shifts in cell type or subset frequencies are biologically significant. In addition to scCODA, we employed the Mann-Whitney U-test and Fischer’s exact test. Differentially abundant cell types and subset had to be identified as significant by at least two of the aforementioned methods.

### Differential expression analysis

Pairwise (i.e., naïve vs IgG; αCD4 vs IgG) differential expression (DE) analyses were conducted using MAST, on log2(TP10K+1) normalized gene expression data (Finak et al., 2015; Kotliar et al., 2020); **Figures 4**, **5**, **6**). The covariates mitochondrial reads and number of genes we included when performing DE.

### Pseudobulk differential expression analysis

To robustly identify DE genes among cell subsets, we performed pseudobulk DE analysis. For cell subsets of interest, we generated pseudobulk counts from scRNA-seq gene expression matrices. Psuedobulk counts and associated metadata (e.g., sample, NHP identity) were imported into R and subject to DE analysis using the DESeq2 package. DE was performed using the Wald statistical test and highlighted genes where selected using the threshold pvalue<0.05 and log_2_(|fold change|)>0.3785.

### Differential cell-cell and receptor-ligand analyses

To discern putative differential cell-cell interactions from our scRNA-seq dataset, we adopted MultiNicheNet. Distinct from conventional interaction cell-cell interaction methods, MultiNicheNet can identify differential, context-dependent cellular communications, leveraging ’pseudobulk’ profiles from scRNA-seq data.

Using MultiNicheNet, we assessed the interaction strength between cell types and identified putative differential cell-cell and ligand-receptor (L-R) pairs – derived from MultiNicheNet’s 50 top-prioritized links (i.e., top 50 predictions across contrasts, senders, and receivers). To highlight highly interconnected cellular populations, we focused on the top 10 (five per experimental group) – as identified in MultiNicheNet’s 50 top-prioritized links – differential interactions per experimental group. To identify cellular subsets underlying differential coarse-level cell-cell L-R among reinfection granulomas (IgG vs naïve, and IgG vs αCD4), we removed all nonimmune cell subsets to identify putative “senders” and “receiver” subsets modulating immune tone in the reinfection granuloma. The top 50 prioritized links among all *IL10*+ T, NK cell sender subsets were queried to identify putative receivers (among all immune cell subsets). The same strategy was employed in determining putative receivers of neutrophil- and monocyte-senders. Interaction matrices were visualized in Python.

## SUPPLEMENTAL FIGURE LEGENDS

**Figure S1.**
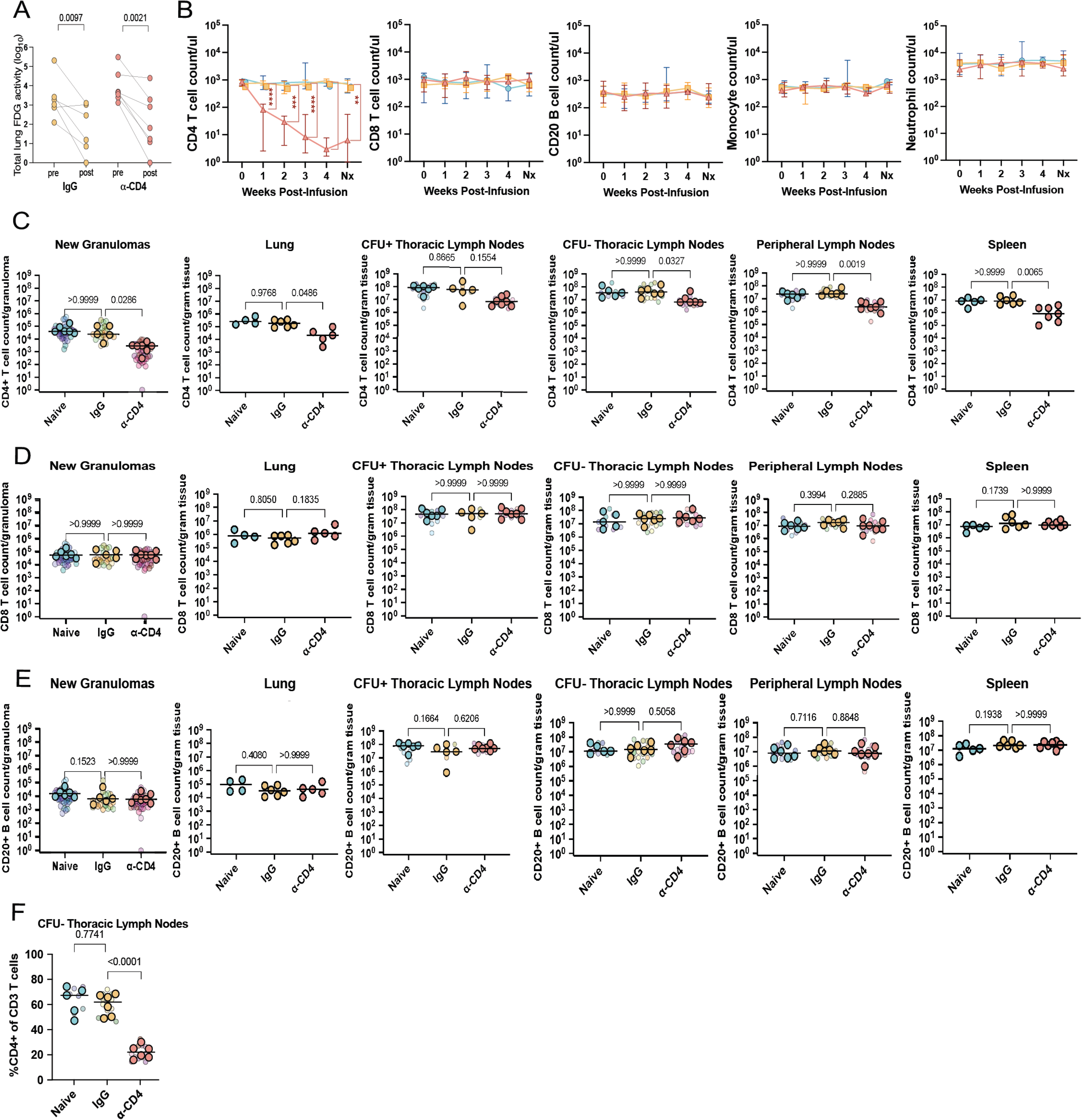
Cellular frequencies across anatomic compartments, related to Figure 1. **(A)** Total lung FDG activity pre- and post-HRZE drug treatment. Two-way ANOVA with Bonferroni’s multiple comparisons test. Dots with connected lines represent individual animals. **(B)** Absolute counts (count/uL) of CD4^+^ T cells, CD8^+^ T cells, CD20^+^ B cells, monocytes, and neutrophils post-antibody infusion in peripheral blood. Median and range shown; mixed-effects model with Dunnett’s multiple comparisons test (** p<0.01, ***, p<0.001, **** p<0.0001). **(C)** Fraction of CD3^+^, CD4^+^ cells from peripheral lymph nodes resected pre- and post-Ab infusion. Dots with connected lines represent individual animals. Wilcoxon matched-pairs signed rank test. **(D)** Absolute counts (count/μL) of CD4^+^ T cells in new granuloma, uninvolved lung, CFU^+^ and CFU^-^ thoracic lymph nodes, peripheral lymph nodes, and spleen. **(E)** Absolute counts (count/μL) of CD8^+^ T cells in new granuloma, uninvolved lung, CFU^+^ and CFU^-^ thoracic lymph nodes, peripheral lymph nodes, and spleen. **(F)** Absolute counts (count/μL) of CD20^+^ B cells in new granuloma, uninvolved lung, CFU^+^ and CFU^-^ thoracic lymph nodes, peripheral lymph nodes, and spleen. **(G)** Fraction of CD3^+^, CD4^+^ cells from CFU^-^ thoracic lymph nodes resected at necropsy. One-way ANOVA with Dunnett’s multiple comparisons test. (D-F) Transparent smaller dots represent granulomas, colored by animal. Larger dots represent mean per animal and lines represent medians. Kruskal-Wallis with Dunn’s multiple comparisons test adjusted p-values reported.

**Figure S2.**
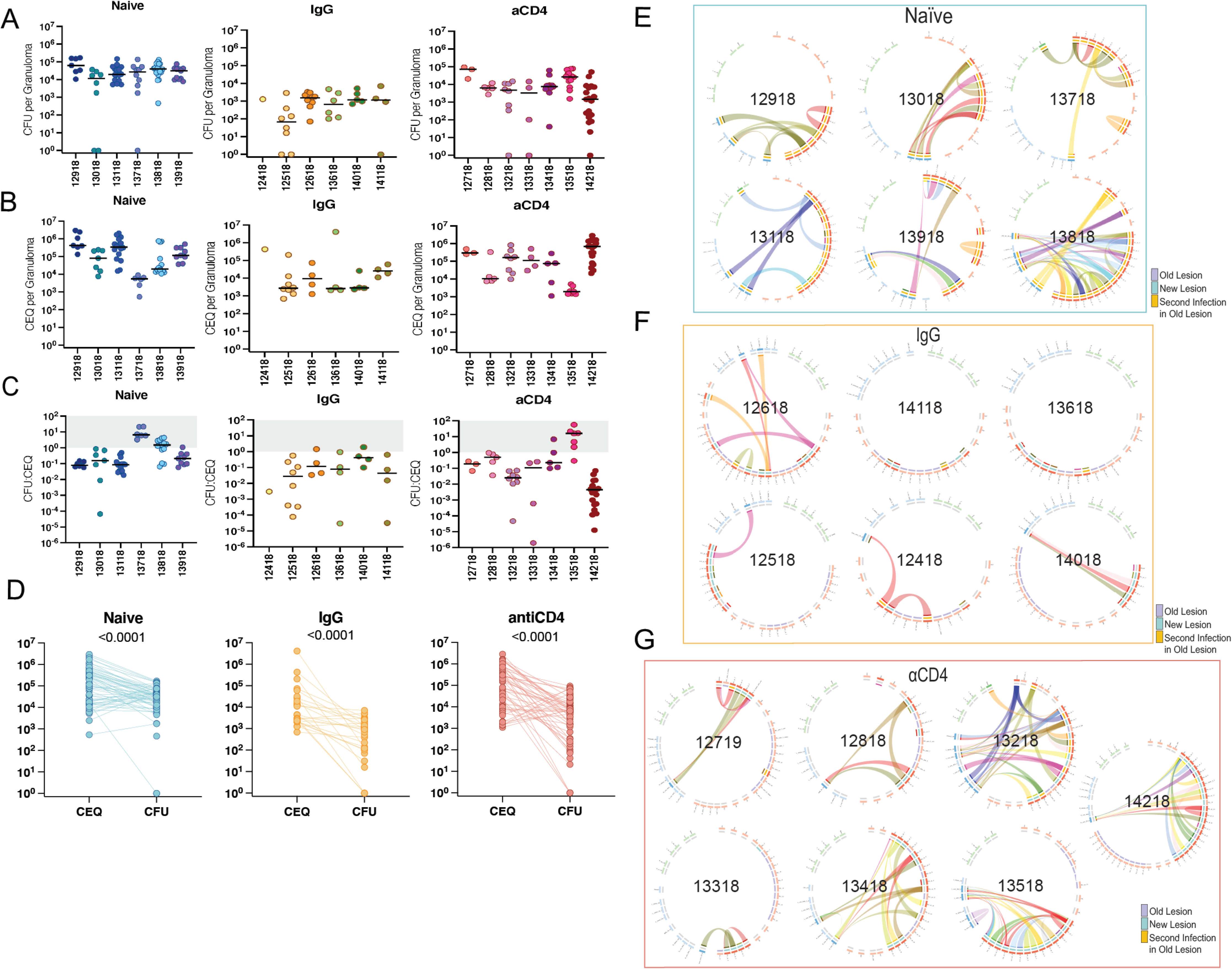
*Mtb* CFU, CEQ, CFU:CEQ, and dissemination, related to Figure 2. **(A)** CEQ (left), and CFU (right). Individual dots represent individual TB granulomas. Linkages depict CEQ and CFU within the same lesion. Wilcoxon matched-pairs signed rank test. **(B)** *Mtb* CFU per granuloma. Integers on the x-axis represent an individual macaque. Colored dots are individual granuloma. Lines represent median per animal. **(C)** *Mtb* CEQ per granuloma. Integers on the x-axis represent an individual macaque. Colored dots are individual granuloma. **(D)** CFU:CEQ per granuloma. Integers on the x-axis represent an individual macaque. Colored dots are individual granuloma. Lines represent median per animal. **(E-G)** Circos plots depicting *Mtb* strains shared between anatomical sites in naïve (E), reinfected (F) or reinfected with CD4 depletion (G) animals. Individual circos plots represent a single macaque, labeled with their study ID. In the plots, each wedge represents a distinct tissue site that was sampled and/or sequenced for barcodes. There are three tracks in each plot. The outer ring defines tissue samples as being from lungs (red), thoracic lymph nodes (blue) or distal extrapulmonary sites (green). Lighter shades represent tissues that were plated for *Mtb* but were sterile. The middle ring represents lesions that were detected by PET-CT during primary infection with the first *Mtb* library (‘old lesion’), new lesions detected by PET-CT after re-infection (‘new lesion’), or new lesions (as defined by Library S barcode sequencing) that were also found at sites where old lesions had previously formed (detected by PET-CT). The innermost ring represents distinct barcodes found in each tissue, where each unique barcoded *Mtb* strain (from the secondary *Mtb* library) is given a different color. Tissues that share the same *Mtb* strain by sequencing are linked by ribbons. Lung tissues are further grouped by lobe, abbreviated as follows: RUL, right upper lobe; RML, right middle lobe; RLL, right lower lobe; LUL, left upper lobe; LML, left middle lobe; LLL, left lower lobe; Acc, accessory lobe.

**Figure S3.**
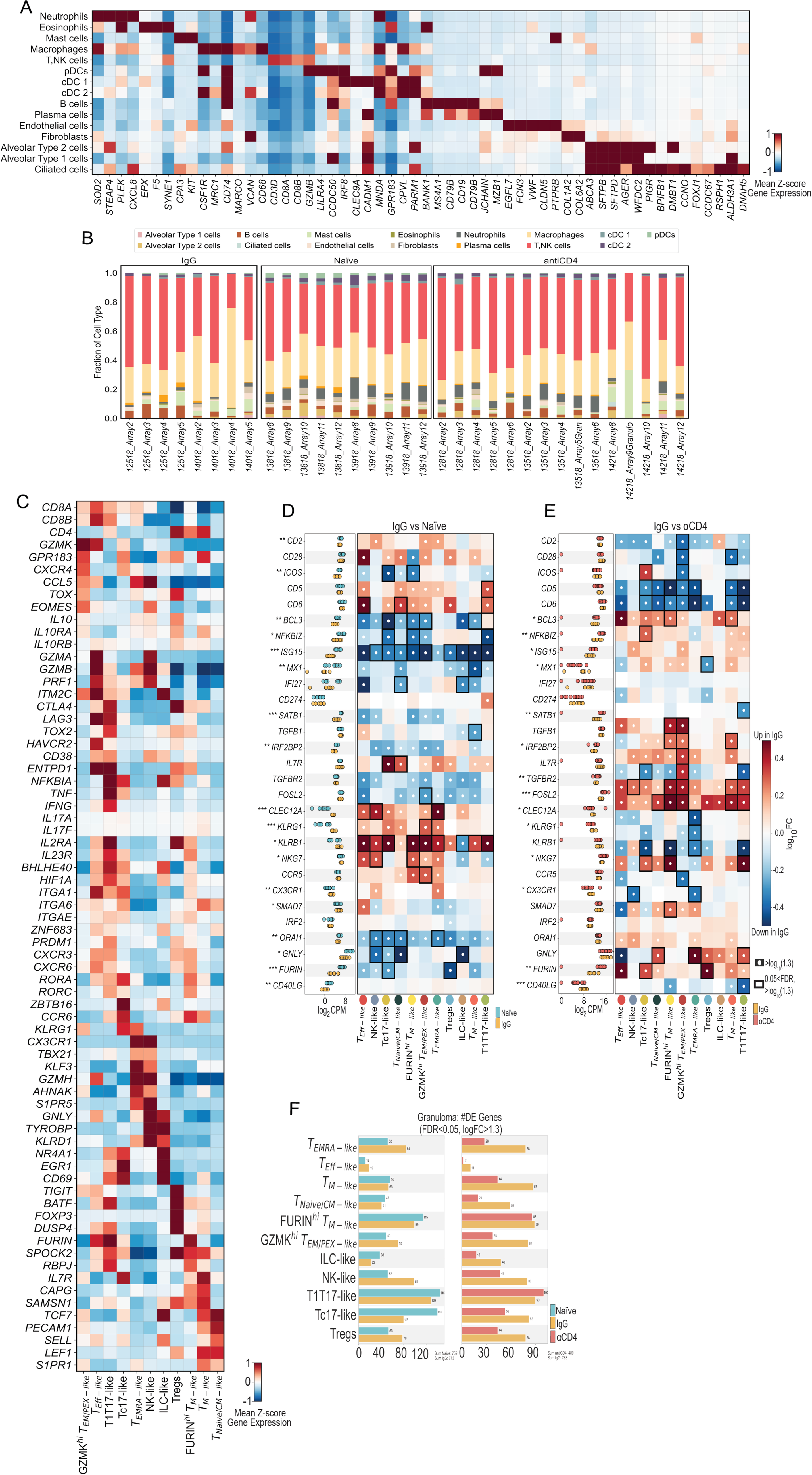
Coarse-level cell types and T, NK cell diversity in the TB granuloma, related to Figures 3, 4. **(A)** Heatmap depicting gene expression profiles (mean z-score) of coarse cell-type markers. Marker genes are depicted on the x-axis, and coarse-cell types on the y-axis. **(B)** Coarse-level cell type frequencies in the TB granuloma (naïve (left), IgG (middle), αCD4 (right)). Individual bars represent a single TB granuloma profiled using Seq-Well S^3^. **(C)** Heatmap depicting gene expression profiles (mean z-score) of T,NK subpopulation markers, as well as select transcription factors, cytokine and chemokine receptors, co-inhibitory and co-activation markers. (**D)** T, NK cell pseudobulk Log_2_CPM for naïve (light blue), IgG (yellow), and αCD4 (red) NHP granulomas (*** p<0.001,** p<0.01, *p<0.05; Wilcoxon rank-sum test). Heatmap depicting log_10_FC of lineage markers, cytolytic molecules, select transcription factors, immunoregulatory molecules, and chemokines, and cytokines (rows) for each cell type (columns). White circles indicate >log_10_|1.3| fold change, relative to naïve or αCD4 granulomas. Black rectangles indicate 0.05 FDR and >log_10_|1.3| fold change, relative to naïve or αCD4 granulomas. **(F)** Number of differentially expressed genes among T, NK cell subpopulations with 0.05 FDR< and >log_10_|1.3| fold change, naïve (light blue), IgG (yellow), and αCD4 (red).

**Figure S4.**
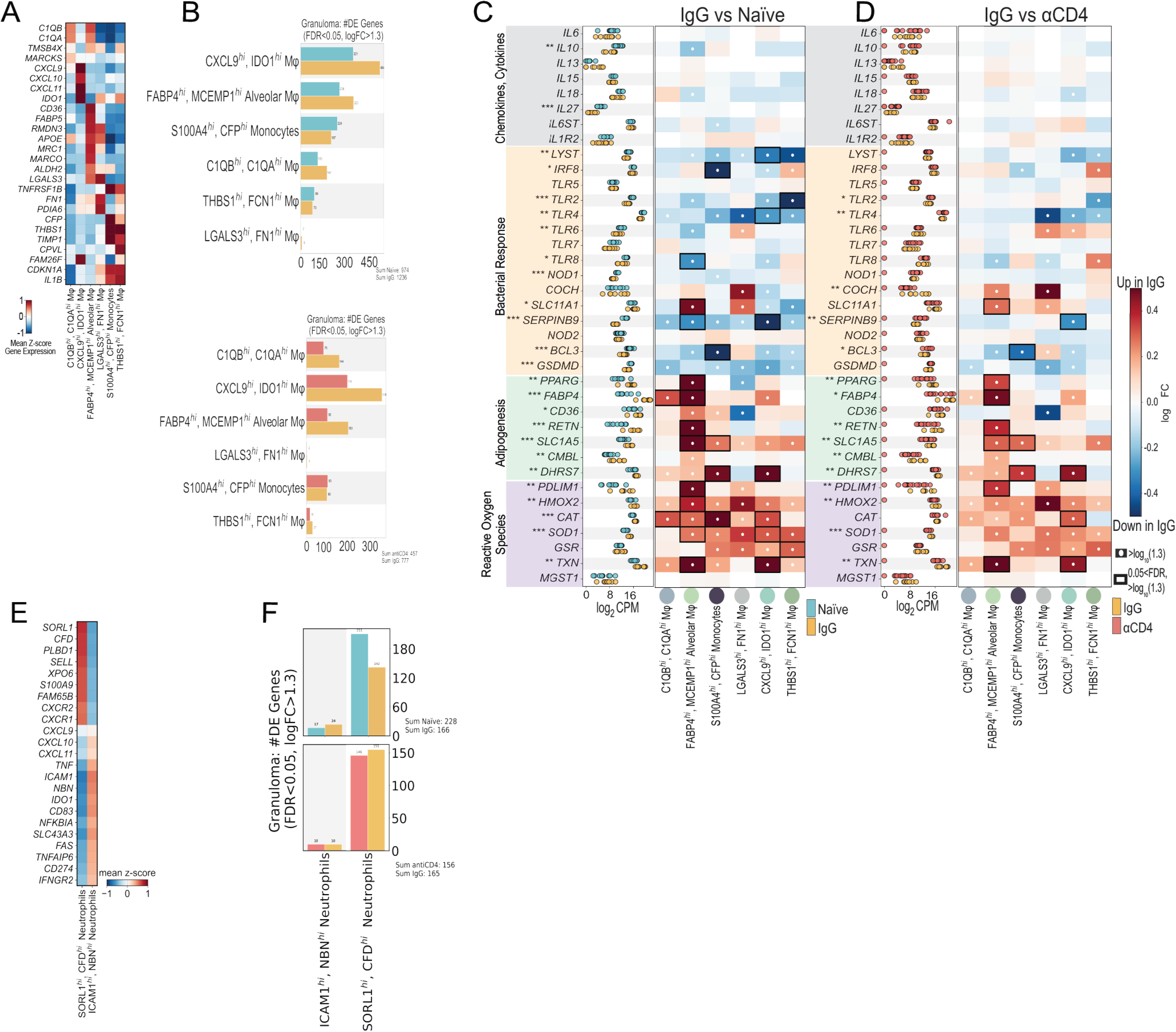
Monocyte-derived gene programming and neutrophil heterogeneity, related to Figures 5, 6. **(A)** Heatmap depicting gene expression profiles (mean z-score) of monocyte-derived subpopulation markers. **(B)** Number of differentially expressed genes among monocyte-derived subpopulations with 0.05 FDR< and >log_10_|1.3| fold change, naïve (light blue), IgG (yellow), and αCD4 (red). **(C-D)** Monocyte-derived pseudobulk Log_2_CPM for naïve (light blue), IgG (yellow), and αCD4 (red) NHP granulomas (*** p<0.001,** p<0.01, *p<0.05; Wilcoxon rank-sum test). Heatmap depicting log_10_FC of select chemokines and cytokines, bacterial response genes (Yao et al., 2018), adipogenesis, and reactive oxygen species (rows) for each cell type (columns) in NHP granulomas, IgG vs naïve (C) or IgG vs αCD4 lesions (D). White circles indicate >log_10_|1.3| fold change, relative to naïve or αCD4 granulomas. Black rectangles indicate 0.05 FDR and >log_10_|1.3| fold change, relative to naïve or αCD4 granulomas. **(E)** Heatmap depicting neutrophil subpopulation marker gene expression profiles (mean z-score). **(F)** Number of differentially expressed genes among neutrophil subpopulations with 0.05 FDR< and >log_10_|1.3| fold change, naïve (light blue), IgG (yellow), and αCD4 (red).

**Figure S5.**
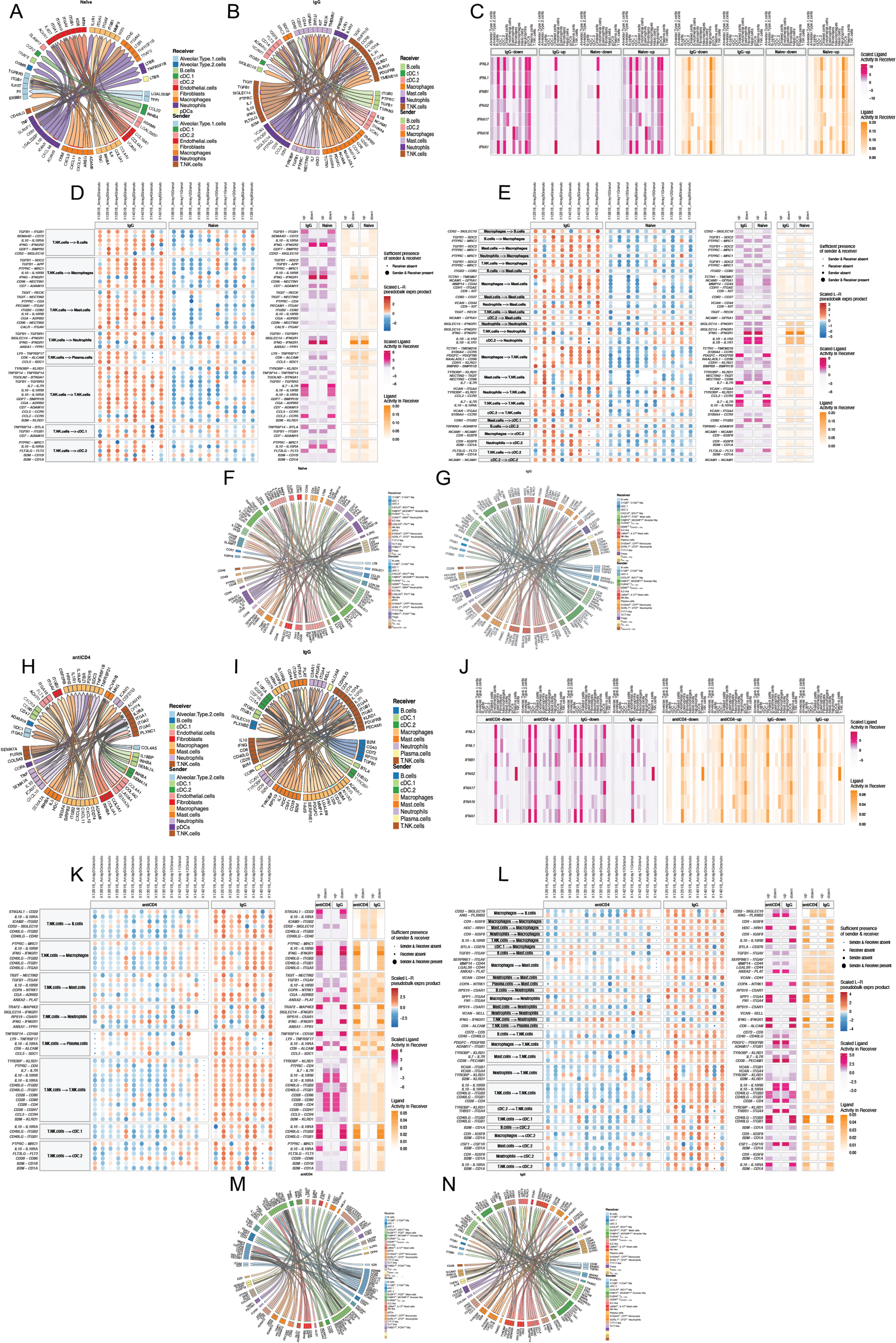
Cell-cell and ligand-receptor interactions during *Mtb* reinfection, related to Figure 7. **(A)** Circos plot depicting differential network interactions among coarse-level cell type annotations in naïve granulomas (naïve vs IgG). Ribbon arrows indicate directionality (sender to receiver) populations. The outer edge color among senders denotes the sender cell type, with the inner edge color representing the receiver cell type. Circos plot depicts the 50 top-prioritized linkages. **(B)** Differential cell-cell interactions among IgG lesions (IgG vs naïve). **(C)** Heatmap depicting type 1 interferon ligand-activity among coarse cell types. **(D)** Heatmap (left) depiction of scaled ligand-receptor expression among the top 50 prioritized linkages among T, NK cells from IgG granulomas, relative to naïve. Columns represent individual granulomas. Heatmap split by NHP group. Heatmaps (right) are representations of the scaled ligand activity among putative receiver populations. **(E)** Heatmap similar to (D) but depicting the top 50 prioritized linkages among all coarse-level cell types. **(F)** Circos plot showing differential network interactions among immune cell subpopulations in naïve granulomas (naïve vs IgG). Top 100 prioritized linkages are plotted. **(G)** Differential cell-cell interactions among all immune cell subpopulations IgG vs naïve. **(H)** Differential cell-cell interactions among αCD4 lesions (αCD4 vs IgG). **(I)** Differential cell-cell interactions among IgG lesions (IgG vs αCD4). **(J)** Heatmap depicting type 1 interferon ligand-activity among coarse cell types, IgG vs αCD4. **(K)** Heatmap similar to (D) but displaying the top 50 prioritized linkages among T, NK cells from IgG granulomas, relative to αCD4. **(L)** Heatmap similar to (D) but depicting the top 50 prioritized linkages among all coarsely annotated cell types from IgG granulomas relative to αCD4. **(M)** Circos plot depicting differential network interactions among immune cell subpopulations in αCD4 granulomas (αCD4vs IgG). Top 100 prioritized linkages are plotted. **(N)** Differential cell-cell interactions among all immune cell subpopulations IgG vs αCD4.

